# Individualized Mapping of Functional Brain Networks in Older Adulthood

**DOI:** 10.64898/2026.01.30.702883

**Authors:** Colleen Hughes, Anne C. Krendl, Roberto C. French, Shannon L. Risacher, Yu-Chien Wu, Andrew Saykin, Richard Betzel

## Abstract

The functional network architecture of the aging brain undergoes significant systematic and idiosyncratic changes. Emergent individualized network mapping approaches may yield better or more sensitive explanatory insight about age-related neural and behavioral variability, although most applications have focused on young adults. In the current study, we tested the validity and impact of mapping individual-specific topography in two fMRI datasets comprising 112 young (18-35 years) and 176 older adults (60-92 years). Older adults had more idiosyncratic network topography than young adults. Individualized maps from resting-state fMRI improved network homogeneity and fidelity to task fMRI activations, while also exhibiting intra-individual reliability and inter-individual discriminability over a 2-year interval. Last, traditional group-averaged (*vs*. individualized) network mapping had a moderate-to-large impact on individual-level estimates of network segregation, a widely-studied measure of functional brain aging. Therefore, individualized network mapping captures important heterogeneity in older adulthood and may yield more precise characterization of neurocognitive aging.

## Background

### Individualized Mapping of Functional Brain Networks in Older Adulthood

A large proportion of the population in many countries now and in the coming decades is entering older adulthood (e.g., 1 in 6 people in the U.S.A. in 2020 ^1^). For this reason, there is a critical need to understand how aging affects people’s health and daily life to promote longer periods of well-being. Studies of brain aging reveal marked differences across the adult lifespan which, in turn, relate to cognitive decline, the quality of social relationships, and symptomatology in forms of non-normative aging such as dementia ^2–5^. A major unifying framework for quantifying brain aging is the organization of brain regions into networks ^6^. Brain network architecture revealed by functional connectivity (FC; i.e., inter-regional co-activations during functional magnetic resonance imaging [fMRI]) increases predictive accuracy of individual differences in cognition and behavior compared to anatomical and structural features^7^. This advantage of FC may be particularly useful for understanding the greater neural and behavioral inter-individual variability observed in older adulthood ^8,9^. That said, most studies rely on group-representative networks – whose topographical maps (i.e., spatial extent and boundaries) are determined based on averages of hundreds or even thousands of brains ^10^ – to calculate and summarize interregional FC. Emergent research contradicts the idea that group-averaged network maps are a good representation of individuals ^11–15^. But, few studies have examined this issue among older adults who are broadly characterized by their heterogeneity ^9^ and for whom ignoring this heterogeneity may have pernicious consequences (e.g., poorer sensitivity to cognitive decline).

Capturing individual-specific network topography in older adulthood is vital for a number of reasons (for a review, see Perez et al. ^9^). For instance, areal boundaries based on FC are weaker and more variable in older age, such that even group-averaged network maps made solely on older adults are a poorer fit to a given older individual than a young adult group map would be to a young individual ^16^. Age group differences may be exacerbated artificially as a result; an interpretation supported by a recent study showing that group-averaged network maps overestimated the magnitude of differences in FC between individuals with schizophrenia compared to a control group ^17^. Also, a preponderance of studies have identified age-related differences in functional activations when participants actively engage in tasks ^18–20^ and rely on spatial maps of networks for contextualizing and interpreting such findings (and *vice versa*). For example, neural activations are less sensitive to increasing cognitive task difficulty in older (*vs*. young) adults, which was attributed to age differences in network organization ^21^. Yet, recent studies in young and middle-aged adults using individualized network maps find less widespread evidence of overlapping functional neuroanatomy (e.g., for different executive functions) ^22,23^.

That is, functional localization of different cognitive functions within individuals is poorly captured by group-level network maps ^22^. Greater fidelity to individual-specific network borders in older adults may thus reveal why some older adults exhibit more (*vs*. less) cognitive impairment. In sum, studies of neurocognitive aging will gain better or unique explanatory insight about heterogeneity in older adulthood from individualized network mapping, which may improve clinical translation.

Template matching is one widely-deployed approach to generate individualized network maps that has made important basic science and clinical contributions ^12,14,24–26^. This approach uses priors to map well-established functional brain networks ^27^ in individuals. Importantly, it can be reliably implemented on shorter fMRI acquisitions ^13^ that are common to extant longitudinal studies examining within-person change during aging in richly phenotyped older adults (e.g., ^28^). Such datasets provide a wealth of information with which to understand the sources of age-related inter-individual variability in brain and behavior in older adulthood. Few investigations, however, have comprehensively established the validity of individualized mapping when applied to aging populations and how individual variation in network topography differs by age ^9^ – which was the central aim of the current study. Across two datasets, we therefore compared individualized *versus* traditional group-averaged network mapping on several evaluation metrics such as network homogeneity, intra-participant reliability, inter-participant discriminability, and network-function alignment. We also characterized the extent of individual and age-related variation in network topography, and moreover, how individualization affected a widely-studied FC summary metric linked to higher risk of cognitive impairment in older adulthood ^2^. By conducting these tests, our objective was to highlight the benefits of individualized network mapping for studying neurocognitive aging and indicate specific challenges to doing so in developmental and clinical populations.

## Results

In the current work, we analyzed two datasets. After data quality exclusions, the first dataset comprised 77 older adults (61-92 years) and 112 young adults (18-35 years) recruited as part of a larger study on normative social cognitive aging at Indiana University (IU) Bloomington. Participants underwent 15 minutes of resting-state, 15 minutes of passive movie-watching, and the false belief task (a localizer for social cognition) ^29^ during fMRI on a Siemens 3.0T Prisma Fit MRI scanner. The second dataset comprised 99 cognitively normal adults (60-88 years) recruited and diagnosed at the Indiana Alzheimer’s Disease Research Center (IADRC).

The IADRC cohort served to replicate and extend observations in the IU older adult cohort. IADRC participants underwent 10 minutes of resting-state fMRI on a Siemens 3.0T Prisma MRI scanner. For participant demographics, see Table 1. For a summary of the participant workflow across analyses, see Supplemental Figure 1. For MRI acquisition and preprocessing details, see Methods.

**Table 1.** Sample description.

|  | **IU**<br>**Young Adults** | **IU**<br>**Older Adults** | **IADRC**<br>**Cohort** |
| --- | --- | --- | --- |
|  | *M (SD) / n* | *M (SD) / n* | *M (SD) / n* |
| **Sample size** | 112 | 77 | 99 |
| **Age in years** | 21.89 (4.17) | 72.88 (6.23) | 71.33 (5.92) |
| **Age in years (min-max)** | 18-35 | 61-92 | 60-88 <sup>a</sup> |
| **Sex (F/M/Non-binary)** | 69 / 40 / 3 | 52 / 25 / 0 | 73 / 26 / 0 |
| **Race** |  |  |  |
| White | 80 | 75 | 68 |
| Black | 2 | 0 | 30 |
| Asian | 20 | 2 | -- |
| Multiracial | 10 | 0 | -- |
| Other | -- | -- | 1 |
| **MoCA** | 27.61 (1.90) | 26.60 (2.48) | 25.72 (2.78) |
| **Education <sup>b</sup>** |  |  |  |
| High school or less | 20 | 1 | 11 |
| Some college / no degree | 56 | 12 | 20 |
| College degree | 21 | 21 | 54 |
| Advanced degree | 15 | 41 | 14 |
| **% Censored Frames** |  |  |  |
| Rest | 8.82 (9.17) | 13.74 (10.03) | 3.57 (4.43) |
| Movie Run 1 <sup>b</sup> | 7.13 (8.59) | 10.17 (8.74) | -- |
| Movie Run 2 <sup>b</sup> | 6.73 (8.97) | 9.74 (8.47) | -- |
| False Belief Task Run 1 <sup>b, c</sup> | 0.77 (1.79) | 2.68 (3.27) | -- |
| False Belief Task Run 2 <sup>b, c</sup> | 1.23 (2.59) | 2.17 (2.91) | -- |

*Note.* Montreal Cognitive Assessment (MoCA) <sup>30</sup>. <sup>a</sup> Exact visit age in years was unavailable for 2 participants. <sup>b</sup> Education level was unavailable for two IU older adults. <sup>b</sup> Only some participants qualified for analyses using these scans – see Supplemental Figure 1 and the Methods (*image quality assessment*). <sup>c</sup> While rest and movie-watching were censored using a 0.1 mm filtered framewise displacement threshold, the false belief task was censored using a different threshold of 0.9 mm framewise displacement appropriate for GLM estimation of task fMRI contrasts <sup>31</sup>.

### Individualized network mapping approach

In the current report, we used a variant of the template matching approach to estimate individual-specific network topography of 14 well-characterized functional brain networks ^12,13,15,32^. In brief, grayordinates for each participant were assigned a single network label based on the similarity of their activation timeseries to spatial priors (i.e., templates) of the 14 networks, which were derived from an independent sample (www.midbatlas.io) ^15^. We combined template matching with a sub-sampling procedure to ensure that individualized network maps were obtained using equal amounts of low motion data, even though older adult participants had more in-scanner movement (Table 1). For detailed procedures, see the *Methods*. Examples of the resulting individualized network maps are shown in Figure 1a and Supplemental Figures 2-3.

**Figure 1.**
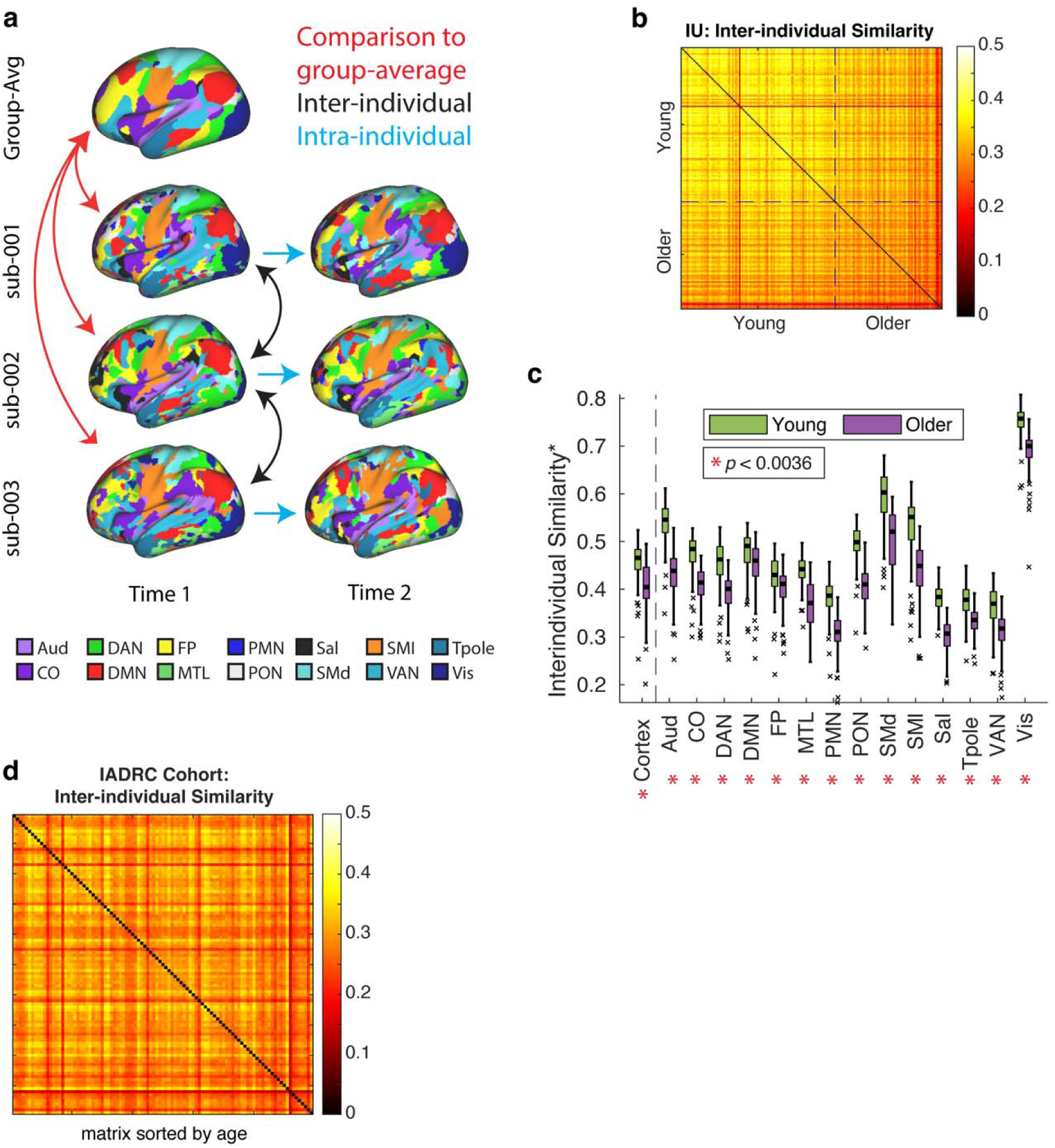
Older adults are more idiosyncratic than young adults. *Note*. (a) Schematic of the analytic approach. We performed three types of comparisons. We compared group-averaged network maps to individualized maps (red arrows); individualized maps from one participant to those of another (black arrows); individualized maps from a single participant at time 1 with maps from the same participant at time 2 (blue arrows). (b) IU cohort (Older: n=77; Young: n=112): A participant × participant matrix indicating inter-individual similarity (normalized mutual information) across cortex where rows/columns were sorted by age in years to show how the oldest older adults have the least inter-individual similarity to their peers. On-diagonal elements (i.e., self-comparison) are not applicable. (c) IU cohort: Boxplots indicating age group differences in inter-individual similarity for cortex and, to the right of the dashed line, each individual network (*dice coefficient for binary vectors). Age differences in inter-individual similarity at the Bonferroni-corrected *p*-value of 0.0036 are noted with an asterisk. (d) IADRC cohort (n=99): Replication of panel b that older age related to less inter-individual similarity.

### Analyses focused on cortical maps

#### Older adults have more idiosyncratic network maps than young adults

First, we tested the central premise of the current work that network topography of older adults would be more different from the group-averaged map compared to young adults. Because most group-averaged network maps in the literature were not created by sampling older adults (e.g., ^10^; c.f. ^16,33^), which could introduce bias for age group comparison, we created group-averaged maps from subsets of participants in the current datasets. Specifically, for the IU sample, we constructed the group-averaged network mapping by applying an analogous template matching procedure to group-averaged FC from a subset of the 50 lowest motion older adults and 50 motion-equated young adults (Supplemental Figure 4). We also created an IADRC-specific group-averaged network map from a subset of the 50 lowest motion IADRC participants (Supplemental Figure 5).

For each participant in the IU cohort, we quantified the normalized mutual information (NMI; ^34^) – a metric between 0-1 where higher values indicate more similar network assignments on a vertex-to-vertex level – between each individual’s map and the group-averaged network map. As hypothesized, in the IU cohort, older adults (*M* = 0.41, *SD* = 0.05) were less like the group-averaged map than young adults (*M* = 0.46, *SD* = 0.04), *t*(187) = 7.61, *p* < 0.001, *d* = 1.12, 95% CI [0.74, 1.45]. However, because group-averaging induces a smoother (i.e., spatially blurred) and more contiguous network map, we also compared how similar each participant’s network map was to every other participant. Older adults (*M* = 0.28, *SD* = 0.03) were more dissimilar to their peers than young adults (*M* = 0.35, *SD* = 0.03), *t*(187) = 16.83, *p* < 0.001, *d* = 2.48, 95% CI [1.78, 3.02]. Notably, lower chronological age related to higher inter-individual similarity among IU older adults, *r*(75) = −0.55, *p* < 0.001, 95% CI [−0.69, −0.37] (Figure 1b). In the IADRC replication cohort, lower chronological age also related to greater inter-individual similarity, *r*(97) = −0.26, *p* = 0.01, 95% CI [−0.43, −0.06] (Figure 1d). In contrast, IU young adults had less variability in chronological age, and it was unrelated to their inter-individual similarity, *r*(110) = −0.18, *p* = 0.06, 95% CI [−0.35, 0.01].

Both global and network-specific changes in FC are observed in older adulthood ^3,35,36^. As such, we next compared inter-individual similarity of specific networks to test the possibility that some, but not all, networks exhibit greater topographical variation in older adults. Each participant’s individualized network map was a vector of values 1-14 indicating the network assignments of each of 59,412 cortical vertices. For each network, we binarized the vector (1 = target network, 0 = any other network) and compared the binarized vectors using the dice coefficient ^15^. We found that all networks exhibited reduced inter-individual similarity in older *versus* young adults, *t*s > 4.26, *p*s < 0.001 (Figure 1c). Taken together, these results suggest that older adults had greater topographical heterogeneity than young adults, which would otherwise not be captured in a group-averaged network map.

#### Individualized network maps increase network homogeneity

In the previous section, we showed that older adults had more idiosyncratic network topography than young adults. It remained unclear, however, if this finding reflected meaningful variation. To establish the validity of individualized network maps, we examined several metrics (for a review, see Perez et al. ^9^, Li et al. ^37^). First, we computed a measure of network homogeneity by taking the mean product moment correlation between every pair of vertices within the same network (e.g., DMN-DMN but not DMN-VIS), following past work ^13,38^.

Greater values indicated that vertices with similar FC profiles were appropriately grouped together. We compared homogeneity values from the vertex-by-vertex dense FC matrix (59,412 × 59,412 cortical vertices) that was divided into networks based on both an individual’s personalized map as well as the group-averaged map. We hypothesized that we would observe greater network homogeneity for individualized *versus* group-averaged maps. We did so by comparing the observed *t*-value to a permuted null distribution created by randomizing the correspondence of mapping method labels to values across 10,000 iterations. Model *p*-values were considered statistically significant if they fell below a Bonferroni-corrected threshold of *p* = 0.0036 (α = 0.05 divided by 14). As hypothesized, individualized maps had greater homogeneity than the group-averaged maps in both age groups across cortex and in most, but not all (i.e., posterior medial network), networks (Figure 2a, Supplemental Figure 6a, Supplemental Table 1). This finding, including non-significance of the posterior medial network, was replicated in the IADRC cohort (Figure 2c, Supplemental Figure 6b, Supplemental Table 2). Global and network-specific improvements in network homogeneity validated that the template matching approach optimized cohesiveness within distinct networks, while preserving inter-participant network homology (i.e., all networks were represented in all participants, which aids inter-participant comparison).

**Figure 2.**
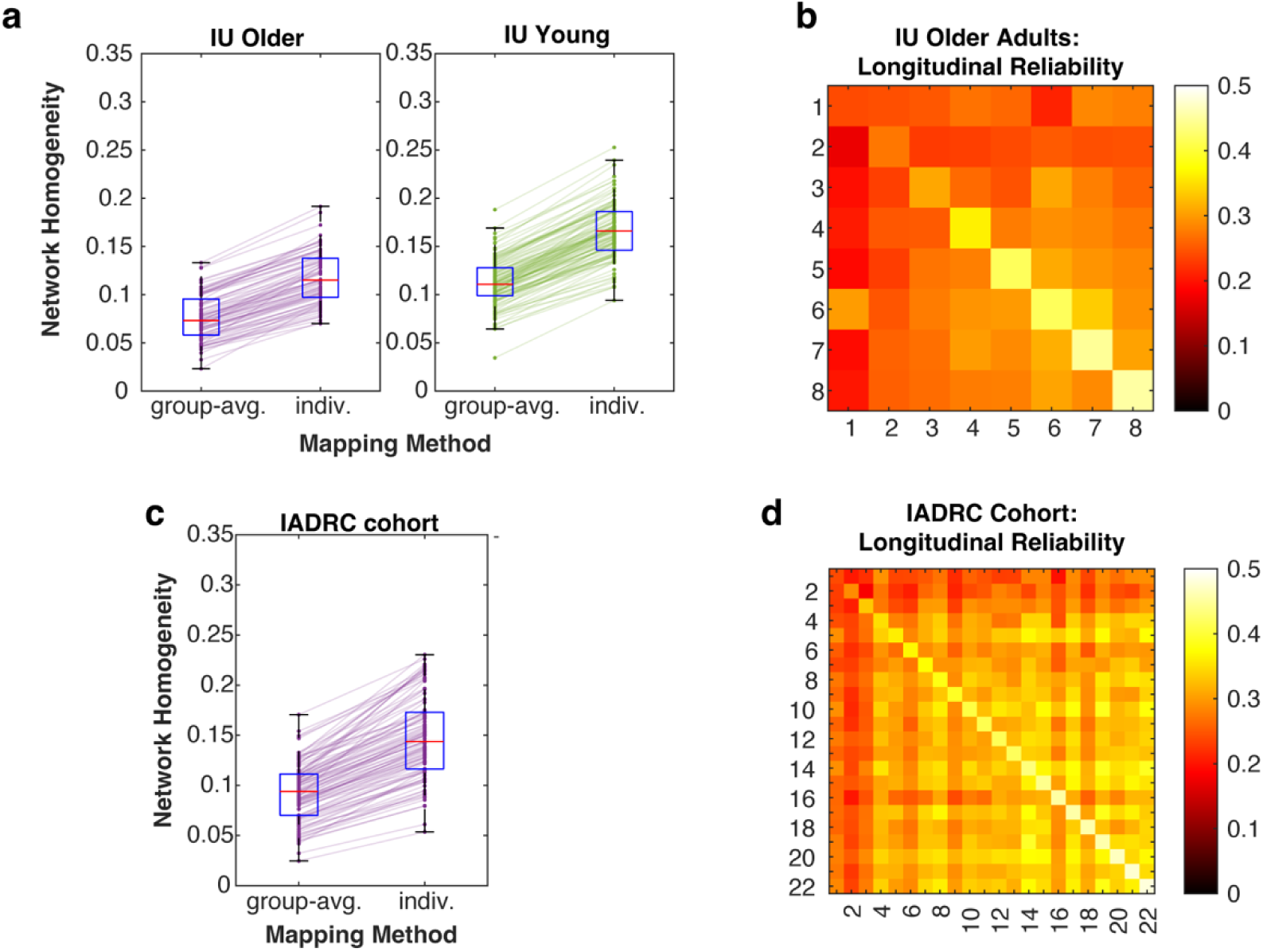
Individualized network maps increased network homogeneity, and intra-individual over a 2-year interval was higher than inter-individual similarity. *Note*. Network homogeneity – the strength of the correlations between vertices within networks – was higher for individualized *versus* group-averaged network mapping in (a) IU older and young adults and (c) the IADRC cohort. Individualized network maps were reliable and discriminable (i.e., more like self than anyone else) over a 2-year interval in a subset of IU cohort older adults (n=8; b) and IADRC cohort older adults (n=22; d) who underwent repeated imaging. The diagonal indicates intra-individual reliability over time. The upper (time 1) and lower (time 2) triangles indicate inter-individual similarity at each time point.

#### Intra-individual reliability is higher than inter-participant similarity across a 2-year interval

Next, we determined whether individualized network maps were reliable within individuals and discriminated between individuals (i.e., individuals were more like their own network maps than anyone else) over time. Although FC weights (i.e., topology) using group-averaged network definition have been shown to be reliable and discriminable within and across testing sessions spanning months to years ^8,33,39^, few studies have examined this question for individual network topography in older adults either cross-sectionally or longitudinally. To address this gap, we leveraged a subset of older adult participants in both the IU and IADRC cohorts who underwent imaging two or more times approximately 2 years apart. We chose this interval because it is common to longitudinal studies of aging (e.g., ^28^). In the IU cohort, 8 older adults (4 male, 4 female, time 1: ages 68-81, *M* = 74.48, *SD* = 4.24) were recruited 2 years (698-768 days) after their first participation to undergo an identical resting-state fMRI protocol.

Individualized network maps were created following the same procedures described above. The maps within individuals over time (intra-individual reliability) and between individuals (inter-individual similarity) were compared using the above-described NMI metric. Specifically, inter-individual similarity was calculated between each pair of participants at each of the two timepoints. Then, we took the average per pair across timepoints. Finally, we took the average across participants in the time-averaged inter-individual matrix, resulting in a 1:1 ratio of intra-individual to inter-individual observations. Statistical significance was determined by permuting values associated with intra-individual and mean inter-individual comparisons.

In the IU older adult longitudinal cohort (n = 8), participants were more like themselves over time (*M* = 0.37, *SD* = 0.08) than like others (*M* = 0.26, *SD* = 0.02), *t*(7) = 3.42, *p* = 0.005, *d* = 1.62, 95% CI [0.54, 3.27] (Figure 2b). In the IADRC cohort (n = 22, 6 male, 16 female, time 2: ages 58-77, *M* = 68.76, *SD* = 4.73) whose imaging sessions were approximately 2 years apart (379-994 days, *M* = 610.55, *SD* = 186.50), we replicated higher intra-individual reliability (*M* = 0.40, *SD* = 0.06) than inter-individual similarity (*M* = 0.30, *SD* = 0.03) over time, *t*(21) = 7.13, *p* < 0.001, *d* = 2.11, 95% CI [1.02, 3.12] (Figure 2d). These findings demonstrate that individualized mapping is a feasible strategy to deploy in longitudinal studies of within-person change. See the Supplemental Material for similar analyses, conducted within a single testing session by pooling rest and passive movie-watching fMRI data^13,40,41^, that compare young *versus* older adults.

#### Individualized network mapping explains more variance in task activation patterns compared to group-averaged network mapping

Findings of overlapping functional neuroanatomy from task fMRI studies inform theoretical models across domains ^22,42,43^. For example, activity in default network regions is elicited by tasks that require participants to think about themselves in the past/future and about other people, which can be taken as evidence that personal experiences facilitate understanding of other people’s thoughts, feelings, and behavior (i.e., theory of mind ^43,44^). But, group-averaged mapping methods may overestimate the shared neural basis of these forms of social cognition ^23^. For this reason, we next tested the alignment between network boundaries and task activations to validate individualized network mappings ^37,40,45^. Specifically, IU young and older adults additionally underwent fMRI while engaging in the well-established false belief task ^46–48^ for localizing neural activations associated with theory of mind. Theory of mind exhibits robust age-related behavioral impairment ^49^ and neural differences in activations and FC ^50–52^, making it well-suited to the current investigation.

To measure network-function alignment for each participant, we followed an established spatial ANOVA approach ^38^ in which activation values for the theory of mind condition > control condition (Figure 3a) *z*-contrast were predicted by network assignment using a one-way ANOVA. Variance explained (*R*^2^) was the outcome of interest, where higher values indicated better network-function alignment. We compared the observed *R*^2^ values to two null models: other participants’ individualized network maps ^38^ (Figure 3b) and the group-averaged network map (Figure 3c), each applied to a given participant’s contrast map. In both cases, we observed that participant’s own individualized network maps explained more task variance in both older adults, *t*s > 3.37, *p*s ≤ 0.001 and young adults, *t*s > 3.95, *p*s < 0.001. The observed greater variance explained in older adults is noteworthy because older adults tend to have weaker theory of mind activations in regions such as medial prefrontal cortex, precuneus, and temporal poles ^29,48^ which could have, but did not, limit the greater precision afforded by individualized network mapping. Gaining individual-level specificity to functional neuroanatomy through individualized mapping may thus promote more precise characterization of behavioral and neural deficits occurring in some, but not all, older adults.

**Figure 3.**
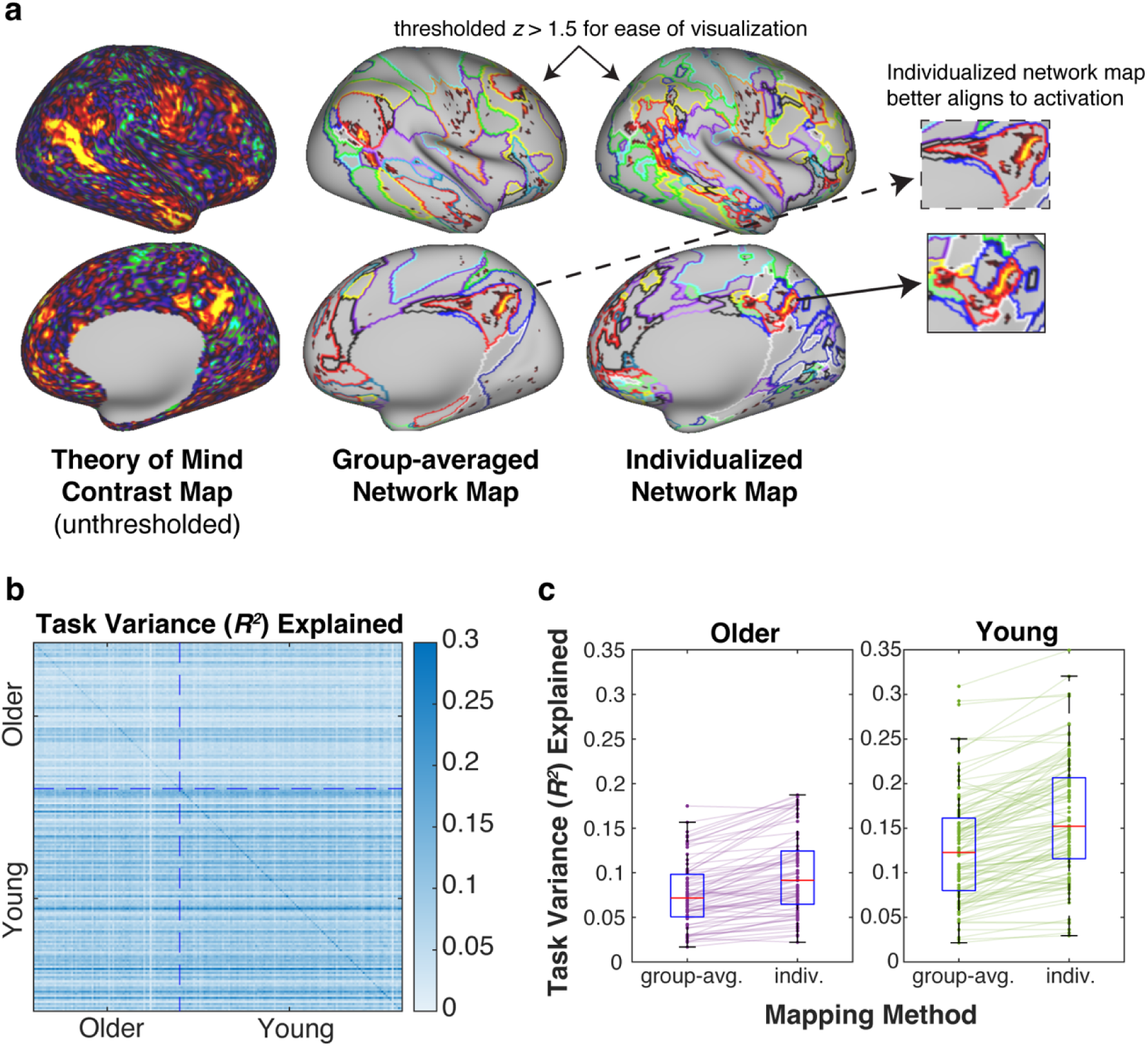
Individualized network mapping improves network-function alignment with theory of mind task fMRI contrast activations in both young and older adults *Note*. To examine network-function alignment, we first calculated the GLM *z*-contrast [theory of mind condition > control condition] for each participant (a, left). Network-function alignment was operationalized as the variance in the task activation map that was explained by the individualized map (a, right; b, on-diagonal). We compared the observed *R^2^* for each participant to two null models: variance explained by other participants’ individualized maps (b, off-diagonal) and the group-averaged network map (a, middle; c). Under both null models, participants’ own individualized networks explained more task activation variance.

#### Network topography

Having validated the homogeneity, reliability, discriminability, and external validity to task activity of individualized network maps in older adults, we next sought to describe age-related topographical similarities and differences that would be otherwise unobservable using group-averaged network maps. We examined three complementary features of age-related network topography: (i) probabilistic similarity, (ii) spatial consensus, and (iii) network size and displacement.

#### Probabilistic network topography is similar across age groups

Prior reports indicated that some areas of cortex have highly consistent network assignments within and across samples ^13,15^. For comparison to this literature, we calculated the probability of each network in each group (i.e., the fraction of participants to which each vertex was assigned to a given network).

Qualitatively, the probabilistic maps were similar across age groups (Supplemental Figure 8). To quantify these observations, we calculated the similarity (i.e., product-moment correlation across all vertices) of the unthresholded probabilistic maps between young and older adults. Indeed, the probabilistic maps between young and older adults were highly correlated across networks (*r*s = 0.96 – 0.99). We observed, however, that older (*vs*. young) adults tended to have less consensus with their peers in both peripheral and some, but not all, central areas of certain networks (e.g., the dorsal anterior cingulate area of the cingulo-opercular network [CO]).

It is common to compare individuals in terms of FC and contextualize those results using network labels. However, these network-level summaries may be misleading because many grayordinates have inconsistent network assignments across individuals. As an alternative, recent work has suggested that these types of analyses may benefit by comparing individuals using only those grayordinates with a high level of consensus in their network assignments, leading to improved reliability of brain-behavior correlations ^13^. This prompts the next question that we addressed: are high consensus areas similar between young and older adults? We identified highly consensual network assignments in each IU age group and the IADRC cohort by taking the modal network assignment at each vertex, calculating the probability of that modal assignment, and thresholding to retain vertices with 80% or greater consensus (Figure 4a). We observed that high consensus areas were broadly similar in location across groups, but the spatial extents (i.e., size) at the 80% threshold tended to be smaller in older adults. This observation was supported by binarizing the thresholded maps to vertices labeled as high consensus or not and comparing their spatial similarity between young and older adults. Young and older adults had lower similarity at the 80% threshold for high consensus regions (dice coefficient = 0.77) than any comparison between 10,000 permutations of random groups, *p* < 0.001. For results at other thresholds, see Supplemental Figures 9-10. The finding that older adults had less consensus than young adults is consistent with the previously reported finding that their individualized network maps were more idiosyncratic.

**Figure 4.**
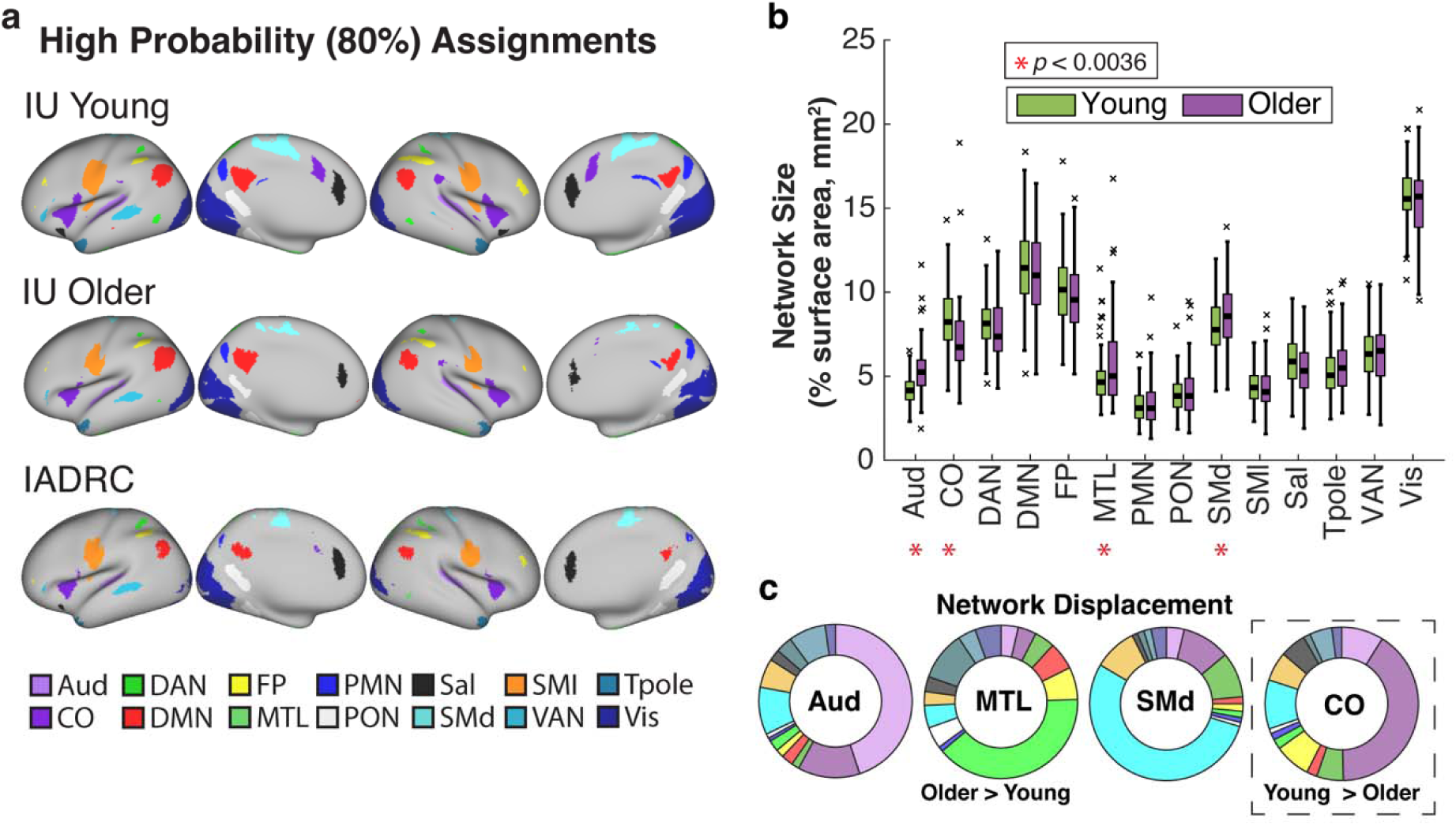
Individualized network topography *Note.* (a) High probability network assignments in each group using an 80% probability threshold (see also Supplemental Figure 9, which uses a 60% threshold). (b) Some networks exhibited significant age differences (IU: young *vs*. older) in surface area, noted with an asterisk if below the Bonferroni-corrected *p*-value of 0.0036. (c) For those networks, the donut plots indicate the average percent of vertices where network assignments agreed between groups or not, and if not, which of the other networks was commonly assigned to the same vertex.

#### Some, but not all, networks differ in size in older *versus* young adults

Based on recent work in individuals with and without depression ^32^, another feature of network topography that could differ by age and is uniquely afforded by individualized, but not group-averaged, mapping is network size. Specifically, we calculated the percent of surface area in mm^2^ of vertices assigned to each network in each person ^32^. Using a Bonferroni-corrected *p*-value threshold of *p* = 0.0036, we found that the following networks were expanded in older *versus* young adults: auditory network, *t*(187) = 6.55, *p* < 0.001; medial temporal lobe network, *t*(187) = 3.14, *p* = 0.001; and dorsal somatomotor network, *t*(187) = 2.94, *p* = 0.0036 (Figure 4b).

Conversely, the cingulo-opercular network was expanded in young *versus* older adults, *t*(187) = 4.60, *p* < 0.001 (Figure 4b). All other networks did not show a significant difference in size by age group, *t*s < 1.84, *p*s > 0.16. Because extant work varies in reporting network size as a function of the percent of surface area or percent of vertices, we calculated both though they were highly correlated across age groups, *r_mean_* = 0.92, *SD* = 0.03. We also conducted a robustness analysis on the residuals of network surface area after regressing out a participant-level measure of topological complexity – used to quantify surface registration quality ^53,54^ – that was higher in older *versus* young adults, *t*(187) = 3.72, *p* < 0.001. The direction and magnitude of age differences in network size residuals were similar, Aud: *t*(187) = 5.88, *p* < 0.001; CO: *t*(187) = 4.02, *p* < 0.001; but the age differences in the medial temporal lobe network, *t*(187) = 2.43, *p* = 0.012, and dorsal somatomotor network, *t*(187) = 2.36, *p* = 0.009, were no longer significant at the Bonferroni-corrected threshold.

To further characterize these age differences in network size, we conducted an exploratory analysis in which we examined which networks were ‘displaced’ when older adults had expanded auditory, dorsal somatomotor, and medial temporal lobe networks compared to young adults and, conversely, when young adults had an expanded cingulo-opercular network compared to older adults. To do so, we compared network assignments for each vertex between each pair of individuals. Of the vertices where the assignments disagreed, we calculated the percent of vertices attributed to the displaced network. Then, to characterize group-level findings, we averaged the displacement matrix (networks × networks) for older adults compared to young adults and vice versa. This approach mirrors past reports, except it compares individuals to each other rather than to a group-averaged network mapping ^24,32^. Qualitatively, we observed that older adults’ larger auditory network may “trade-off” (exchange border vertices) with the spatially adjacent cingulo-opercular network, which was expanded in young adults (Figure 4c). Future investigations should consider the physiological, neurobiological, or behavioral antecedents and correlates of these differences, as well as potential confounds (e.g., inter-regional temporal signal-to-noise ratio is particularly low in the medial temporal lobe network, Supplemental Figure 11) when interpreting findings about network size (see also ^14,55^).

#### Group-averaged network mapping overestimates age group differences in network segregation compared to individualized network mapping

Topography and topology are related, raising the possibility that individualized topography leads to different estimates of age differences in topology. As such, we next tested how the effect of individualized *versus* group network topography impacted FC analyses.

Specifically, we tested the impact of mapping method on estimates of network segregation, a ratio of within-network FC (homogeneity) to between-network FC (distinctiveness) ^36^. We focused on this FC metric because less network segregation relates to: (1) older age in both cross-sectional ^35,36^ and longitudinal ^56^ studies, (2) less neural activation selectivity ^57^ and less efficient network reconfiguration ^52^ when participants engage in cognition ^58^, (3) cognitive deficit and reserve ^36,56,59^, (4) greater dementia severity ^4^, and (5) greater tauopathy in Alzheimer’s disease ^60^. Altogether, these findings highlight that network segregation is a robust indicator of neurocognitive aging that captures important heterogeneity among older adults.

First, we observed that cortical segregation was greater when using individualized *versus* group-averaged network maps in both the IU, *t*(188) = 5.34, *p* < 0.001 (Figure 5a), and IADRC (Figure 5b), *t*(97) = 2.47 *p* = 0.012, cohorts. This finding reflects that individualized mapping increases both network homogeneity (stronger within-network FC) and network distinctiveness (weaker between-network FC). Also, we replicated past observations that young adults had higher network segregation than older adults, using both the group-averaged, *t*(187) = 5.29, *p* < 0.001, *d* = 0.78, 95% CI [0.49, 1.09]; and individualized, *t*(187) = 4.61, *p* < 0.001, *d* = 0.68, 95% CI [0.37, 0.96] mapping. We then tested the possibility, consistent with our hypothesis and prior work (e.g., Levi et al. ^17^), that group-averaged (*vs*. individualized) network maps overestimate the magnitude of cohort differences. We tested this possibility with a linear model that included age group, mapping method, and their interaction. We extracted the observed *t*-value for the interaction term. Then, we randomly permuted the mapping method labels to create a null distribution of magnitudes of age differences in segregation due to chance. We found that the group-averaged network mapping method yielded a small but significantly greater magnitude of age group differences in segregation compared to individualized network mapping, *t*(1, 378) = 2.20, *p* = 0.043, η_p_ ^2^ = 0.013, 95% CI [0.000, 0.048]. Speaking to how segregation might be used to estimate individual risk, we also tested whether older adults had a greater discrepancy in segregation values between mapping methods than young adults. To do so, we created a participant-level difference score in segregation (individualized minus group-averaged), then compared the observed *t*-value between age groups to a null distribution from permuted age group labels. Older adults showed a greater discrepancy between mapping methods than young adults, *t*(187) = 4.90, *p* < 0.001, *d* = 0.72, 95% CI [0.41, 0.98]; Figure 5 highlights that the individuals in the lowest quartile for group-averaged segregation values had the most discrepancy between mapping methods.

**Figure 5.**
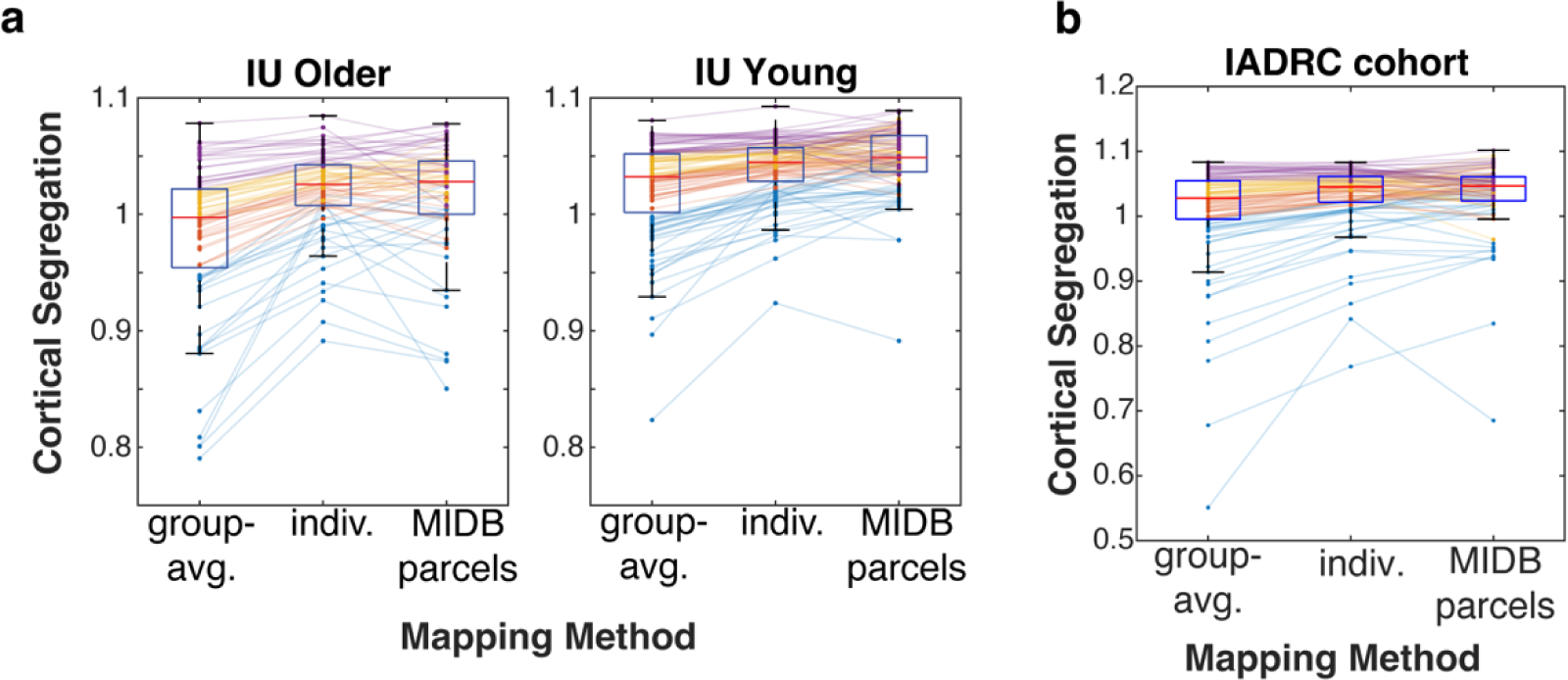
Cortical segregation values by cohort and mapping method (group-averaged network mapping, individualized network mapping, MIDB probabilistic parcellation) *Note.* For the group-averaged (“group-avg.”) and individualized (“indiv.”) mapping methods, segregation was calculated on the dense FC matrix (59,412 cortical vertices). For the MIDB parcellation, segregation was calculated on the 80-parcel FC matrix. Lines represent individual participants and are colored by quartile of segregation values using the group-averaged map. (a) IU older and young adult cohorts. (b) IADRC cohort.

Last, we examined how areal parcellation that leverages, rather than dilutes, individual variation in topography – by excluding vertices whose network assignments are less consistent across individuals– affects network segregation. For this comparison, we used Masonic Institute for the Developing Brain (MIDB; www.midbatlas.io) probabilistic parcels to capture only the regions of cortex whose network assignments are highly consensual across individuals and samples (analogous to Figure 3a) ^13^. Using these 80 regions (14 networks, 2-14 parcels/network), we constructed parcellated FC matrices (80 × 80) for each participant upon which network segregation was calculated. We replicated that young adults had higher network segregation than older adults using the probabilistic parcels, *t*(187) = 5.66, *p* < 0.001, *d* = 0.83, 95% CI [0.50, 1.11]. Unlike the comparison between the group-averaged and individualized methods, there was no evidence that probabilistic mapping affected the magnitude of cohort comparison compared to individualized mapping, *t*(1, 378) = 1.37, *p* = 0.21, η_p_ ^2^ = 0.005, 95% CI [0.000, 0.031].

Nonetheless, older adults again showed a greater discrepancy between mapping methods (individualized *vs*. probabilistic parcels) compared to young adults, *t*(187) = 2.82, *p* = 0.005, *d* = 0.42, 95% CI [0.13, 0.69] (Figure 5a); which suggests that vertices with low consensus convey individual-specific FC patterns. In sum, network mapping that respects, *versus* ignores, inter-individual topographical variation yielded more conservative differences attributable to age and thus may provide more precise estimates of individual-level risk using topological metrics like segregation, thereby improving clinical translation.^26^

## Discussion

An interesting tension exists between the idea of canonical functional brain networks that are observable in individuals ^12^, emerge early in neurodevelopment ^61,62^, and exhibit correspondence across mental states and modalities ^41,63^ but also have substantial inter-individual variation in spatial localization ^13,15^. This tension reflects that functional brain networks are fundamental units by which to understand brain organization across and within individuals.

Network and areal boundaries tend to exhibit greater uncertainty, especially in older age in the context of cerebrovascular and anatomical differences (e.g., cortical atrophy; see network size robustness tests) ^16^, but could also reflect accumulated experience and plasticity ^64^.

Individualized network mapping is therefore a powerful tool for characterizing neural and behavioral heterogeneity that is greater in older adulthood ^9^. To this point, we made several important discoveries. First, network topography in older adults was not only less like widely used group-averaged network maps, but moreover, more idiosyncratic than network topography in young adults. Age-related heterogeneity thus inflated age cohort and individual-level differences in network segregation, a widely studied measure of neurocognitive aging. Second, individualized network maps showed greater consistency within individuals over time than they do between individuals, indicating that even standard acquisition parameters (i.e., in datasets that prioritize large cohorts) capture meaningful inter-individual variation. Third, areas of high network assignment consensus are generally similar across age groups, but these areas represent only approximately half of the cortex, and some networks are more consistent across individuals (e.g., visual) than others (e.g., frontoparietal). Typical approaches assume 100% shared topography, which is why these results – consistent with prior work ^13,15^ – strikingly demonstrate not only the extent of inter-individual but also age-related variation in network topography.

Our finding that older adults are heterogeneous aligns with past work using FC weights (assuming shared topography) ^8,52,65,66^, as well as behavior, anatomy, and clinical presentation ^67–72^. One source of greater variability is older chronological age, as we showed across two samples comprising 176 older adults. Even so, the correlations were moderate, which suggests that other factors contribute to heterogeneity in this period of development. Prioritizing more precise measurements of brain networks moves the field beyond age cohort comparisons (c.f., ^3,16^ a comprehensive assessment of age group differences using an alternate approaches for individualizing regions *vs*. networks), which may better leverage the rich behavioral and neural phenotyping of older adults in extant datasets. For instance, not accounting for topographical variation may explain why there is mixed evidence for prediction of individual differences in cognition, which exhibits age- and disease-related deficits ^73^, from intra-individual (e.g., run-to-run reliability) *versus* inter-individual (e.g., similarity to high performers) features derived from FC weights ^52,74–76^. In fact, we show improvements in network-function alignment even among older adults who, as a group, exhibit weaker neural activations and robust behavioral impairments related to theory of mind ^29,48–50^. The major implication of this finding is that individualized mapping could be used to more precisely identify neural correlates underlying different symptomology in non-normative aging (e.g., episodic memory in Alzheimer’s disease versus impulsivity in frontotemporal dementia). But also, our findings reveal that segregation, a summary metric of functional brain health more broadly, is more precise at the group- and individual-level using individualized mapping. Unacknowledged heterogeneity in functional neuroanatomy limits clinical translation of fMRI-based measures ^26^ and reduces the reliability of brain-behavior correlations ^13^ – altogether supporting the promise of individualization to promote better understanding of older adulthood.

Another key contribution of this work was that individualized mapping yielded reliable and discriminable network topography in two samples of older adults under common acquisition parameters (e.g., <20 minutes of total fMRI at 3.0T using single-echo EPI) that aim to not overburden participants who may find the scanner environment uncomfortable. Importantly, we showed that these individualized maps were reliable and discriminable over a 2-year interval common to gold standard longitudinal studies of within-person aging (e.g., ^28^). This is important because few studies have evaluated the reliability of network topography in any population across such a duration (see also ^32^), making it a relevant observation for aging and non-aging researchers alike. One limitation of this work is that we excluded a relatively large proportion of participants due to conservative motion thresholds, albeit comparable to a prior report using adolescents (i.e., ∼40% of adolescents in the ABCD cohorts in ^13^; 28% of IU older adults). Such challenges of individualized network mapping in traditional acquisitions highlight exciting new directions for precision fMRI ^64,77^. For instance, further analyses reported in the Supplemental Material propose that age differences in topographical reliability could be ameliorated by using different (*vs*. fixed) amounts of data in each group by pooling rest and movie-watching fMRI data ^13,40^, and future work in older adults using longer acquisitions spread out over multiple sessions/days ^33^ may determine optimal ratios. Also, template matching for individualized maps of functional brain networks is a common downstream tool to achieve greater precision, although most work using this approach has largely investigated group-averaged FC templates as opposed to the spatial probability templates used here ^13,15,32^. Precision fMRI in developmental and clinical populations is an active area of inquiry ^3,9,26,33,64,78^; the current work complements the diversity of approaches to gaining precision to individuals and, thus, serves as an important foundation for future insights.

Beyond showing that individualized mapping is valid for studying neurocognitive aging, we also highlight how network topography itself can be a fruitful area of inquiry. Indeed, multiple investigations in young and middle aged adults, adolescents, and clinical conditions have been conducted in recent years ^13,14,24,32,33,55,78^. We conducted an exploratory analysis of age differences in network size given that a recent report differentiated groups of individuals with major depressive disorder from non-clinical controls on the basis of a larger salience network ^24,32^, indicating that network size differences may be a novel risk factor for certain conditions.

Taking a similar approach, we observed that network size differences between age groups appeared to be driven by trade-offs between spatially adjacent networks – termed border shifts ^14,63^. Interestingly, the cingulo-opercular network was expanded in young adults relative to older adults, which corresponded with the near absence of high-consensus cingulo-opercular vertices in dorsal anterior cingulate cortex in older adults; an area spatially adjacent to the dorsal somatomotor network with greater surface area in older adults. Put simply, the cingulo-opercular network was displaced by the dorsal somatomotor network in older *versus* young adults. One reason these networks may be showing age differences is that a more fine-grained network topography – discoverable with precision fMRI ^38,79^ – exists. More work is needed to understand why differences emerge at the group- and individual-level, and the heterogeneity observed in aging is very well-suited for such investigations.

In sum, individualized mapping of functional brain networks in older adulthood is not only feasible but vital for both group and individual-level analyses of network topography and topology. Ignoring inter-individual variation in topography disproportionately mischaracterizes older adults, which we benchmarked to traditional group-averaged mappings across several metrics. That said, we also showed that individualized maps are less reliable in older than in young adults (Supplemental Material) even while evincing other positive characteristics like intra-individual reliability over time and increased network homogeneity. As precision functional mapping methods proliferate, it is critical to test them on developmental and clinical populations to identify and ameliorate persistent biases that would prevent clinical translation in the groups for whom such applications are most valuable.

## Methods

### IU Bloomington young and older adults: Participant information and image acquisition Participants

At Indiana University (IU) Bloomington, young and older adult participants were recruited as part of a larger study on social cognitive aging. Older adult participants were pre-screened for cognitive impairment upon recruitment via a telephone-based, well-validated six-item screener ^80^. A subset of participants without contraindications (e.g., claustrophobia) underwent neuroimaging. Data collection was approved by the Indiana University Institutional Review Board (#11801), including written informed consent from each participant. For the current analyses, participants were included if they underwent neuroimaging (108 older adults, 123 young adults). Of that subset, participants were excluded if they did not complete the resting-state scan or if it had poor image quality (e.g., due to high motion – see *Image quality assessment*; 31 older, 11 young). The analyzed sample comprised 112 young adults (18-35 years) and 77 older adults (61-92 years); see Table 1 for sample demographics. Older adults (range: 19-30) had lower Montreal Cognitive Assessment scores ^30^ compared to young adults (range: 22-30), *t*(182) = 3.17, *p* = 0.002, *d* = 0.47, 95% CI [0.16, 0.75].

### Image acquisition

Neuroimaging was performed with a 20-channel head/neck coil on a Siemens 3.0T Prisma Fit MRI scanner at the Indiana University Imaging Research Facility in Bloomington, Indiana. During the 15 minutes resting-state scan, the projector was turned off (no fixation was presented) and participants were instructed to stay awake and keep their eyes open; no other instructions were given. The movies – two episodes of the mockumentary-style television show Nathan for You (each approximately 8 minutes long; for more information, see Hughes et al. ^52^) – were next presented sequentially using MATLAB version 2022b through a Dell laptop running Windows 10 and a projector illuminating a screen that was visible to participants through a mirror attached to the head coil. Then, participants completed two runs (each 5 minutes 20 seconds) of the false belief task, a well-established localizer for neural activity related to theory of mind *versus* non-social narrative comprehension ^23,46,47^, which was administered using E-Prime version 3; see ^48^ for detailed procedural information. In brief, participants read short written vignettes that required them to make inferences about another person’s mental state (theory of mind) or objects (non-social control condition) by judging subsequently presented statements about the vignettes as true or false; responses were made in the scanner using a button box. Participants viewed a total of 12 vignettes for each condition in a randomized order (i.e., event-related design) across two runs.

The T1w anatomical scan was acquired with a high-resolution 3-D magnetization prepared rapid gradient echo sequence (MPRAGE; 160 sagittal slices, 1.0 mm^3^ isotropic voxels, TE = 2.7 ms, TR = 1800 ms, flip angle = 9°, no acceleration). fMRI scans were collected using a gradient-echo echo-planar image (EPI) sequence sensitive to blood oxygen level dependent (BOLD) contrast (T2*; 450 volumes; 54 interleaved slices, 2.5 mm^3^ isotropic voxels, TE = 30 ms, TR = 2000 ms, flip angle = 70°, A/P phase encoding direction, multi-band acceleration factor = 2). Prior to all functional imaging, phase-encoding polarity reversed (PEPOLAR) spin-echo EPI image pairs were acquired for distortion correction (TE = 58 ms, TR = 7700 ms, flip angle = 90°; image and voxel dimensions matched the resting-state parameters).

A subset of older adult participants underwent an identical resting-state fMRI protocol at the same site and scanner approximately two years later. The movie-watching and false belief task scans were not repeated. The image acquisition parameters for the follow up MPRAGE T1w anatomical were the same as above, with the addition of generalized auto-calibrating partially parallel acquisition (GRAPPA) acceleration factor = 2.

### IADRC: Participant information and image acquisition

#### Participants

IADRC participants were recruited as part of the embedded Indiana Memory and Aging Study (IMAS) and a separate study on social connectedness. Longitudinal clinical, neuropsychological, and MRI data collection took place at Indiana University School of Medicine, Indianapolis. Data collection was approved by the Indiana University Institutional Review Board, including written informed consent from each participant. The analyzed sample comprised 99 older adults (60-88+ years; Montreal Cognitive Assessment scores range: 17-30, 2 not available) who were categorized as cognitively normal by the IADRC Clinical Core team of psychiatrists, neurologists, and neuropsychologists. Only MRI visits that occurred within 1 year of a cognitive testing session were analyzed, and the most recent MRI was used unless it was deemed poor quality (e.g., due to motion – see *Image quality assessment*; Supplemental Figure 1).

For longitudinal analysis of topography, a subset of 22 participants (58-77 years) was selected for analysis because they had at least two good quality MRI sessions that were 1-3 years apart. If more than 1 pair of scans met these criteria, we used the most recent scan pair. One participant was 58 at their first visit but over 60 at the follow up visit; the second, more recent scan was used for cross-sectional analyses.

#### Image acquisition

Neuroimaging for the IADRC cohort was performed on a Siemens 3.0T Prisma MRI scanner with a 64-channel head coil at the Indiana University School of Medicine, Indianapolis. During rest, the projector was turned off (no fixation was presented), and participants were instructed to think of nothing and to remain still with eyes closed. The T1w anatomical scan was acquired with a MPRAGE sequence in accordance with the Alzheimer’s Disease Neuroimaging Initiative (http://adni.loni.usc.edu; ADNI2: 220 sagittal slices, 1.1 × 1.1 × 1.2 mm^3^ voxels, GRAPPA acceleration factor of 2; or ADNI3 [n=25, scans after June 2023]: 1 mm^3^ isotropic voxels). Fluid-attenuated inversion recovery (FLAIR) scans were incorporated into the Freesurfer workflow to refine estimation of the pial surface, when available (n=12 without). One run of resting-state fMRI lasting 10 minutes and 7 seconds was collected using a gradient-echo EPI sequence sensitive to BOLD contrast (T2*; 500 volumes, 54 interleaved slices, 2.5 mm^3^ isotropic voxels, TE = 29 ms, TR = 1200 ms, flip angle = 65°, A/P phase encoding direction, multi-band acceleration factor = 3). Prior to resting-state fMRI, PEPOLAR single-echo EPI image pairs matching the resting-state acquisition parameters (excepting TE = 49.8 ms, TR = 1560 ms) were acquired for distortion correction.

#### IU and IADRC: Image preprocessing

Image preprocessing followed the same procedures across the IU and IADRC datasets, except where noted. Initial anatomical and functional image preprocessing was performed using *fMRIPrep* (IU: v23.2.0; IADRC: v24.1.1) ^81^ (RRID: SCR_016216), which is based on Nipype 1.8.6 ^82,83^ (RRID: SCR_002502). In the IU cohort, functional scans across rest, movie-watching, and task states were processed similarly (unless otherwise noted) for straightforward comparison. In the IADRC cohort, the only functional scans that were analyzed were resting-state. In brief, functional images were realigned to correct for motion, underwent slice-timing correction, underwent distortion correction based on two echo-planar imaging references, and were realigned to a corresponding T1w image. The BOLD timeseries were resampled onto the left/right-symmetric template “fsLR” using Connectome Workbench ^84^. Grayordinates files in connectivity informatics technology initiative (CIFTI) format containing 91k samples were also generated with surface data transformed directly to fsLR space and subcortical data transformed to 2mm resolution MNI152NLin6Asym space. All resamplings were performed with a single interpolation step by composing all the pertinent transformations.

#### Denoising rest and movie-watching fMRI

The resting-state and movie-watching scans from both datasets underwent subsequent denoising – appropriate for FC analyses and similar to extant individualized network mapping work ^13,15,32,33,78^ – using the eXtensible Connectivity Pipeline-DCAN (XCP-D, v0.11.1) ^85,86^. Denoising steps included: (1) discarding the first two volumes of each run to account for potential scanner inhomogeneities, (2) nuisance regression of the *fMRIPrep*-calculated confounding timeseries for the three region-wise global signals extracted within the cerebrospinal fluid, white matter, and global masks, six head motion parameters, the temporal derivatives and quadratic terms for those nine terms, linear trend and intercept terms ^86,87^, (3) temporal band-pass filtering (0.009-0.08 Hz), and (4) censoring high-motion outlier time points using filtered framewise displacement (fFD) greater than 0.1 mm ^88^ – a threshold chosen to conservatively mitigate noise while balancing loss of temporal degrees of freedom in developmental samples who exhibit higher motion ^13,33,78^. Last, the denoised data were spatially smoothed with a 6mm full width at half maximum (FWHM; σ = 2.55) geodesic Gaussian kernel using Connectome Workbench ^84^.

#### Image quality assessment

*fMRIPrep* reports were manually inspected. Two IU participants (1 older and 1 young) failed Freesurfer-based segmentation during *fMRIPrep* and were thus excluded from analysis. Participants were further flagged for exclusion if their resting-state scans were incomplete or had 5 or more minutes (of 15 total minutes) of frames deemed high motion outliers (i.e., exceeding the 0.1 mm filtered framewise displacement threshold for censoring) – 10 young and 30 older adults. In the analyzed IU sample, older adults had more censored resting-state frames than young adults, *t*(187) = 3.49, *p* < 0.001, *d* = 0.51, 95% CI [0.23, 0.85]; which is typical in age cohort studies. See Table 1 for descriptive statistics about the percent of censored frames in each group and scan.

In the IADRC cohort, participants were excluded if concurrent visit age and cognitive status (within 1 year of MRI acquisition) were unavailable, their resting-state scans were incomplete, or resting-state scans had 2 or more minutes (of 10 total minutes) of frames deemed high motion (i.e., exceeding the 0.1 mm filtered framewise displacement threshold for censoring). The cutoffs for quantity of censored data differed between the IU and IADRC to account for multiple factors: (1) different total acquisition times, (2) relative proportion of excluded (vs. included) older adult participants due to high motion at a conservative censoring threshold, and (3) findings from a prior study that argue for maximizing data quantity used for network mapping relative to an approximate minimum of 8 minutes of continuously sampled fMRI data (the amount of data used for IADRC network maps) to achieve discriminability (intra-participant reliability higher than inter-participant similarity) ^13^.

#### IU & IADRC datasets: Detailed individualized mapping procedure

As noted in the main text, we generated individualized network maps of 14 established functional brain networks using a template matching approach, implemented in multiple prior studies ^12,13,15,32^. Following ^32^, which included a small subset of older adults, the network templates we used were unthresholded probabilistic spatial templates (i.e., Supplemental Figure 8) derived on an independent sample of 384 Human Connectome Project participants (mean age of 29.3 years) with at least 52 minutes of low motion data (www.midbatlas.io) ^15^. Although this independent sample did not comprise any older adults, probabilistic network templates were chosen because they show good correspondence across a variety of samples and data-driven network assignment methods ^13,15^. Template matching was conducted on the cortical surface and in subcortical volume (i.e., grayordinates). Networks were assigned at the level of each vertex (cortical surface) using a winner-take-all approach ^12^: first, iteratively calculating the similarity of the timeseries of that vertex to the timeseries weighted by the spatial probability of each of the 14 network templates, and second, by assigning the vertex to the network with the greatest similarity ^32^.

We combined template matching with a sub-sampling procedure to ensure that network estimates were obtained using equal amounts of low motion data, even though older adults had more in-scanner movement. Specifically, for each participant in the IU cohort, we randomly sampled 10 minutes of the available low-motion data (total acquisition duration was 15 minutes) and, given these data, performed template matching to obtain an estimate of network assignments. In the IADRC sample, we sampled 8 minutes of the available low motion data; the total acquisition duration was 10 minutes. We repeated this procedure 10 times, as in ^13^, yielding 10 versions of the network maps for each participant. The consensus network label for each grayordinate was its modal assignment across those versions (i.e., the network to which a vertex was most frequently assigned). Last, network clusters smaller than 50 mm^2^ were reassigned by dilating adjacent network clusters using Connectome Workbench ^84^.

#### IU dataset: Theory of mind localizer analysis

The *fMRIPrep* cifti-formatted false belief task outputs were spatially smoothed with a 6mm FWHM (σ = 2.55 sigma) geodesic Gaussian kernel using Connectome Workbench ^84^ and then submitted to task general linear model (GLM) analysis at the individual-level. The data were modeled using *FitLins* (v.0.11.0)^89^, a wrapper for nilearn-based estimation of task GLMs that was developed to take BIDS-style inputs from *fMRIPrep* and analyze them according to a BIDS Stats Model^90^ file to improve the reproducibility of task fMRI analysis. For each participant and run, we modeled the task as a GLM with four conditions reflecting the trial type (theory of mind, control) × trial component (read story, make inference), as well as covariates of no interest from the *fMRIPrep*-derived confounds file: 6 head motion parameters, non-steady state volumes, and volumes exceeding 0.9 mm framewise displacement ^31^. For each participant, we localized neural activity specific to theory of mind using the average theory of mind > control *z*-value contrast images across runs ^23,29^ (Figure 3a, left). Participants were excluded from localizer analysis if either or both task runs were not acquired (e.g., due to time constraints, 4 young, 2 older), or if either run had 25% or more volumes flagged as a high motion outlier (2 young, 4 older).

## Acknowledgements

The authors thank Skylar Wilson, Dr. Clare Johnson, Dr. Lucas Hamilton, Samuel Naranjo Rincon, the Indiana University Bloomington Imaging Research Facility, and the IADRC staff and researchers for assistance with data collection. We also thank the IUB participants, as well as the Indiana Memory and Aging study participants and their family members, without whom this research would not be possible.

## Data availability statement

The IU young and older cohort data supporting the conclusions of the current work are available upon request to the first author. The IADRC cohort data supporting the conclusions of the current work could be requested via https://medicine.iu.edu/research-centers/alzheimers. The Dworetsky HCP probabilistic network templates are publicly available at www.midbatlas.io.

## Code availability statement

The current report used the following public code repositories to generate and analyze features of individualized network maps: https://github.com/DCAN-Labs/compare_matrices_to_assign_networks (Hermosillo et al., 2024, *Nature Neuroscience*) and https://github.com/cjl2007/PFM-Depression (Lynch et al., 2024, *Nature*). Effect sizes and their bootstrapped 95% confidence intervals were calculated using the MATLAB Measures of Effect Size Toolbox (https://github.com/hhentschke/measures-of-effect-size-toolbox).

## Author contributions

**Colleen Hughes**: Data curation, Formal analysis, Investigation, Visualization, Writing—original draft; **Anne C. Krendl:** Conceptualization, Funding acquisition, Investigation, Methodology, Project administration, Supervision, Writing—review & editing; **Roberto C. French**: Data curation, Investigation, Methodology, Writing—review & editing; **Shannon L. Risacher**: Data acquisition, Data curation, writing – review & editing; **Yu-Chien Wu**: Data acquisition, Data curation, Writing – review & editing; **Andrew Saykin**: Funding acquisition, Data acquisition, Writing – review & editing; **Richard Betzel**: Conceptualization, Funding acquisition, Methodology, Supervision, Writing—review & editing.

