## Supplemental Material for "Individualized Mapping of Functional Brain Networks in Older Adulthood"

Colleen Hughes,^1^ Anne C. Krendl,^1^ Roberto C. French,^1^ Shannon L. Risacher,^2,3,4^ Yu-Chien Wu,^3,4^ Andrew Saykin,^3,4^ & Richard Betzel ^5,6^

^1^ Psychological and Brain Sciences Department, Indiana University, Bloomington, IN, USA

^2^ Department of Internal Medicine, Section on Gerontology and Geriatric Medicine, Wake Forest University School of Medicine, Winston-Salem, NC, USA

^3^ Indiana Alzheimer's Disease Research Center, Indiana University School of Medicine, Indianapolis, IN, USA

^4^ Center for Neuroimaging, Indiana University School of Medicine, Indianapolis, IN, USA

^5^ Department of Neuroscience, University of Minnesota, Minneapolis, MN, USA

^6^ Masonic Institute for the Developing Brain, University of Minnesota, Minneapolis, MN, USA

Correspondence should be addressed to:

Colleen Hughes, Ph.D.

Department of Psychological and Brain Sciences

Indiana University

1101 East 10^th^ Street

Bloomington, Indiana USA

**Intra-individual reliability is higher than inter-participant similarity within session**

To examine age differences in intra-session reliability, we pooled rest and passive movie-watching fMRI data acquired within one imaging session in the IU cohort of young and older adults (30 minutes acquisition in total). For a schematic of the acquisition procedure, see Supplemental Figure 7a. After excluding participants who did not complete the movie-watching scans or whose scans had excessive motion, a subset of 69 older adults and 101 young adults had at least 10 minutes each of low motion rest and movie-watching data. Prior investigations showed that maps created from rest and task fMRI are largely consistent and pooling across them is feasible ^22,40,50^. Even so, we created maps using equal number of TRs from each state. We did so because older adults have functional connectivity estimates that are more dissimilar between these states than young adults ^13,49,51^, and it was unclear how much impact having different proportions of each state would have on estimates of age differences in mapping reliability.

To assess reliability, we held out 10 minutes of low motion data that was randomly selected within each task state. We created one individualized mapping on the held-out data. Then, we created additional maps based on 1-10 minutes of the remaining low motion data in 1-minute increments. To determine within-participant reliability, we compared each mapping based on test data to the mapping made on the held-out data. First, we observed that mapping reliability generally increased with more test data, consistent with prior observations ^22^. Second, we observed that older adults had less reliable maps at each time increment compared to young adults, *t*s > 12.36, *p*s < 0.001 – at 10 minutes, *d* = 1.94, 95% CI [1.50, 2.34]. We next asked if we could ameliorate the age difference in reliability by using disparate amounts of low motion data for young *versus* older adults. In the range we tested, we only observed a lack of evidence for an age difference in reliability when creating individualized maps based on 1 minute of low motion data in young adults *versus* 10 minutes of low motion data in older adults, *t*(168) = 0.66, *p* = 0.51, *d* = 0.10, 95% CI [-0.22, 0.39] (Supplemental Figure 7b [annotated]).

**Supplemental Table 1**

*IU cohort: Effects of Mapping Method (Individualized, Group-Averaged) Within and Between Age Groups on Network Homogeneity*

| **Network** | **Mapping Method:**  **Young (n=112)** | | **Mapping Method:**  **Older (n=77)** | | | **Δ-Mapping Method: Young vs. Older** | |
| --- | --- | --- | --- | --- | --- | --- | --- |
|  | *t* | *p* | | *t* | *p* | *t* | *p* |
| Cortex ^a,b,c^ | 14.99 | < 0.001 | | 9.98 | < 0 .001 | 7.93 | < 0.001 |
| Aud ^a,b,c^ | 9.50 | < 0.001 | | 7.47 | < 0 .001 | 3.74 | < 0.001 |
| CO ^a,b,c^ | 11.99 | < 0.001 | | 11.15 | < 0 .001 | 5.35 | < 0.001 |
| DAN ^a,b,c^ | 12.78 | < 0.001 | | 9.66 | < 0 .001 | 6.48 | < 0.001 |
| DMN ^a,b,c^ | 17.79 | < 0.001 | | 13.03 | < 0 .001 | 6.00 | < 0.001 |
| FP ^a,b,c^ | 19.12 | < 0.001 | | 15.47 | < 0 .001 | 7.25 | < 0.001 |
| MTL ^a,b,c^ | 10.64 | < 0.001 | | 8.64 | < 0 .001 | -4.29 | < 0.001 |
| PMN | 2.57 | 0.010 | | 1.08 | 0.287 | 1.83 | 0.069 |
| PON ^a,b^ | 3.66 | < 0.001 | | 4.44 | < 0 .001 | -0.96 | 0.334 |
| SMd ^a,b^ | 4.21 | < 0.001 | | 4.24 | < 0 .001 | -0.94 | 0.346 |
| SMl ^b^ | 2.72 | 0.007 | | 2.94 | 0.003 | -1.35 | 0.175 |
| Sal ^a,b,c^ | 11.77 | < 0.001 | | 11.24 | < 0 .001 | 3.15 | 0.001 |
| Tpole ^a,b^ | 8.86 | < 0.001 | | 9.57 | < 0 .001 | 0.21 | 0.840 |
| VAN ^a,b,c^ | 13.83 | < 0.001 | | 10.96 | < 0 .001 | 3.33 | 0.001 |
| Vis ^a,b,c^ | 7.12 | < 0.001 | | 3.38 | 0 .001 | 5.92 | < 0.001 |

*Note.* The reference level for the Mapping Method factor is individualized. The *p*-values are uncorrected for multiple comparisons; we used a Bonferroni-corrected threshold of *p*=0.0036 to determine statistical significance. Superscripts in the Network column summarize the statistical significance of each term: ^a^ Mapping Method: Young, ^b^ Mapping Method: Older, ^c^ Interaction between Mapping Method (individualized minus group-averaged) and Age Group.

**Supplemental Table 2**

*IADRC cohort (n=99): Effects of Mapping Method (Individualized, Group-Averaged) on Network Homogeneity*

| **Network** | **Mapping Method** | |
| --- | --- | --- |
|  | *t* | *p* |
| Cortex | 10.87 | < 0.001 |
| Aud | 8.17 | < 0.001 |
| CO | 9.63 | < 0.001 |
| DAN | 6.90 | < 0.001 |
| DMN | 13.79 | < 0.001 |
| FP | 12.23 | < 0.001 |
| MTL | 7.55 | < 0.001 |
| PMN | 0.08 | 0.94 |
| PON | 6.43 | < 0.001 |
| SMd | 4.81 | < 0.001 |
| SMl | 3.92 | < 0.001 |
| Sal | 10.26 | < 0.001 |
| Tpole | 6.20 | < 0.001 |
| VAN | 8.22 | < 0.001 |
| Vis | 4.99 | < 0.001 |

*Note.* The reference level for the Mapping Method factor is individualized. The *p*-values are uncorrected for multiple comparisons; we used a Bonferroni-corrected threshold of *p*=0.0036 to determine statistical significance.

**Supplemental Figure 1**

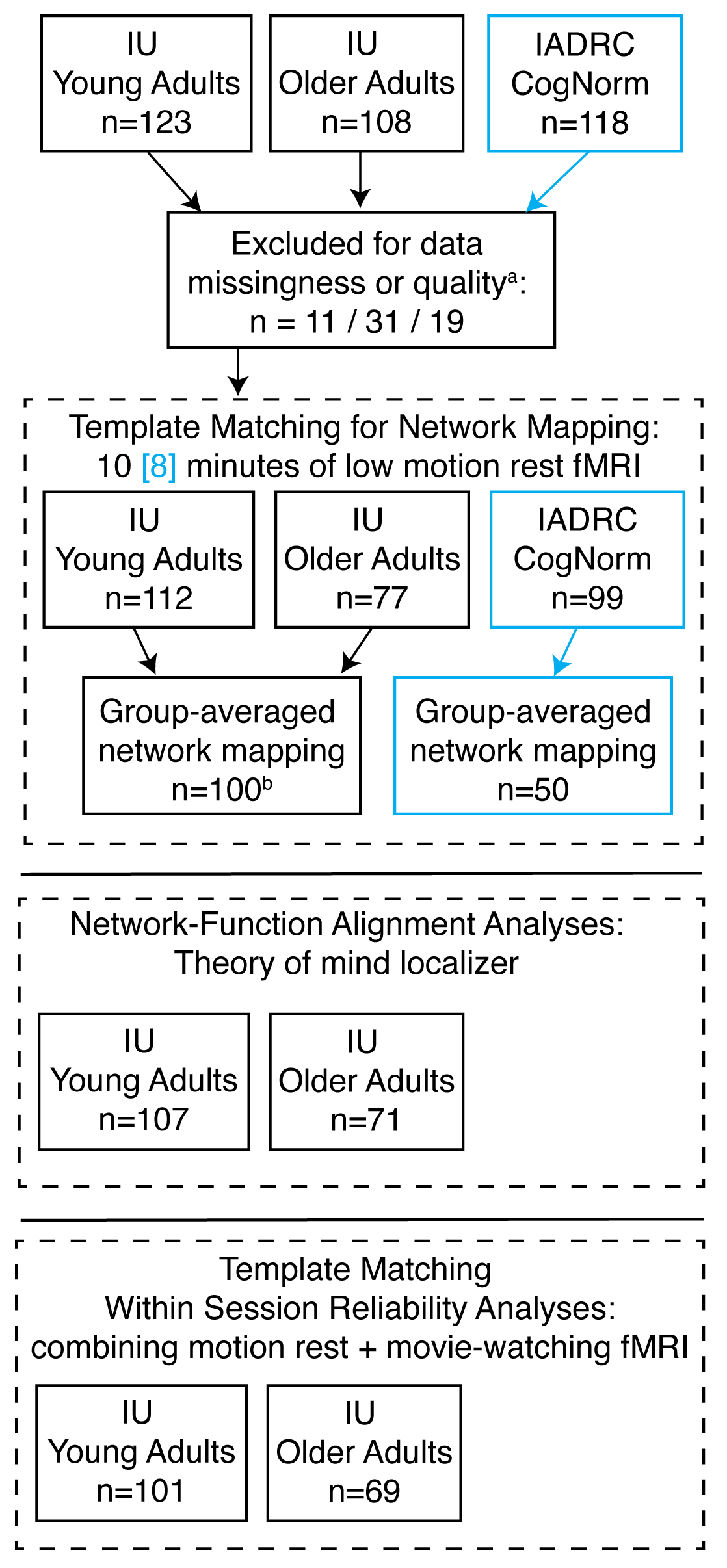
*Participant Inclusion & Workflow*

*Note*. Different subsets of participants were included for different analyses reported across the manuscript. ^a^ See the Methods section *Image quality assessment* for more detail. ^b^ A subset of the 50 lowest motion older adults and motion-matched sample of 50 young adults.

**Supplemental Figure 2**

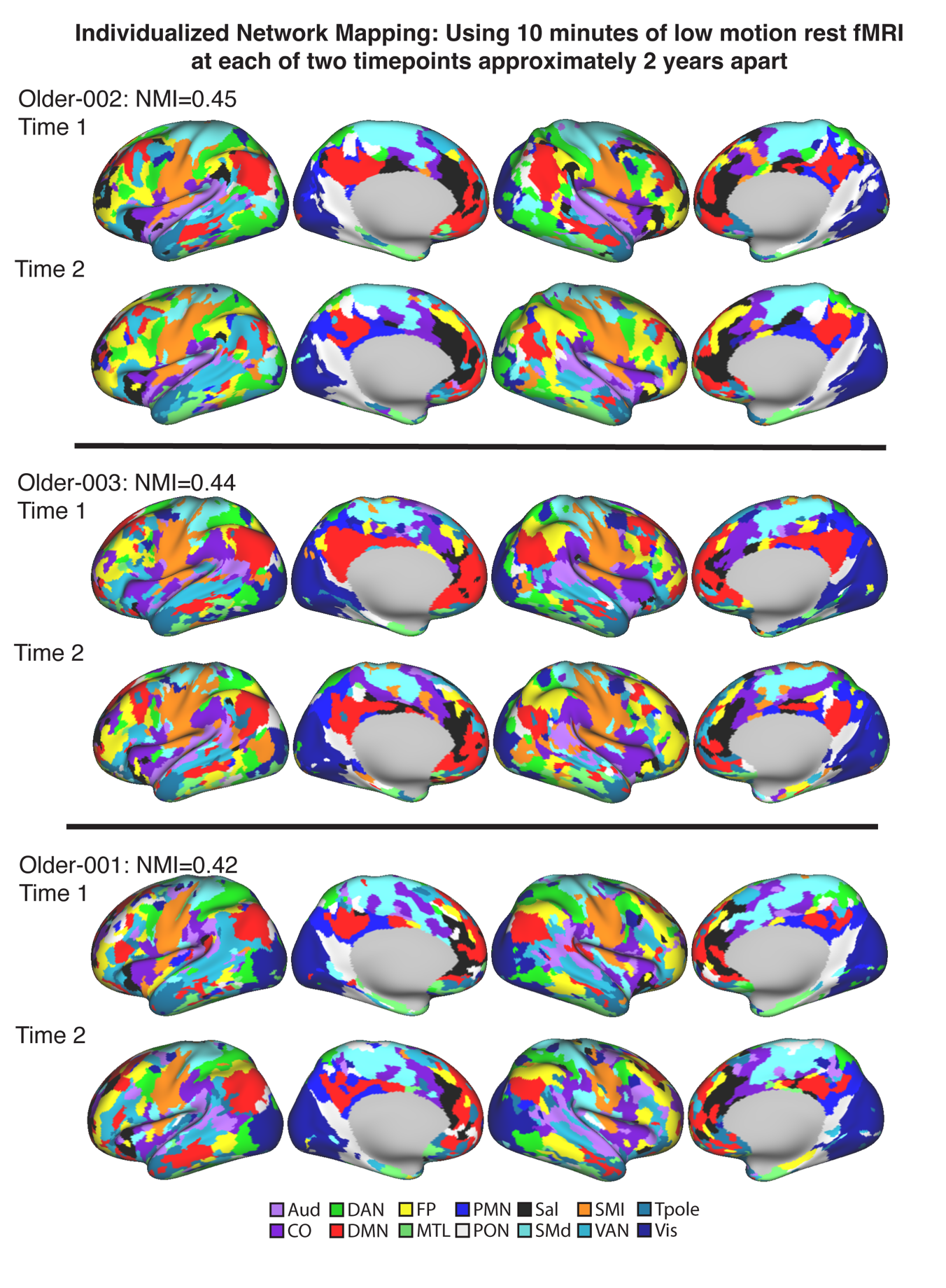
*Exemplar longitudinal individualized network maps in cortex: IU cohort*

**Supplemental Figure 3**

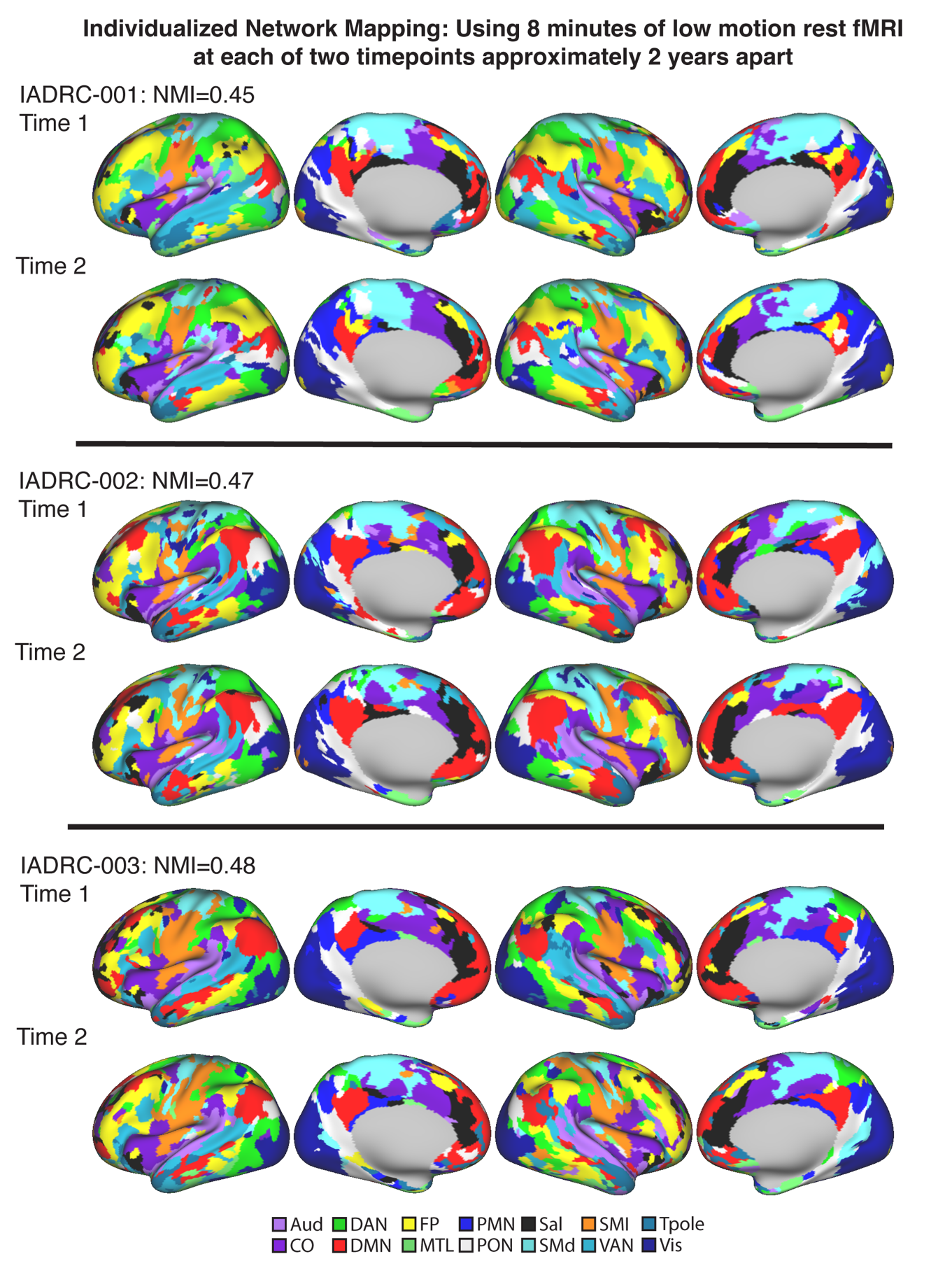
*Exemplar longitudinal individualized network maps in cortex: IADRC cohort*

**Supplemental Figure 4**

*Group-averaged network map: IU subsample*
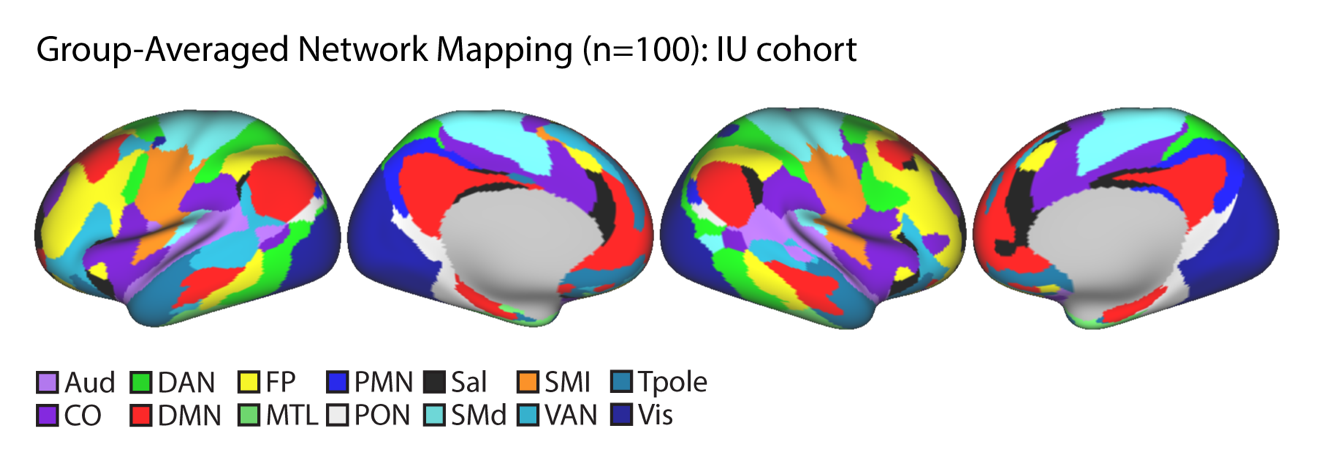

*Note*. This network map was created from the 50 (of 77) lowest motion older adults and a motion-matched sample of 50 (of 112) young adults. It is plotted on the HCP S1200 group-averaged very inflated surfaces.

**Supplemental Figure 5**

*Group-averaged network map: IADRC subsample*

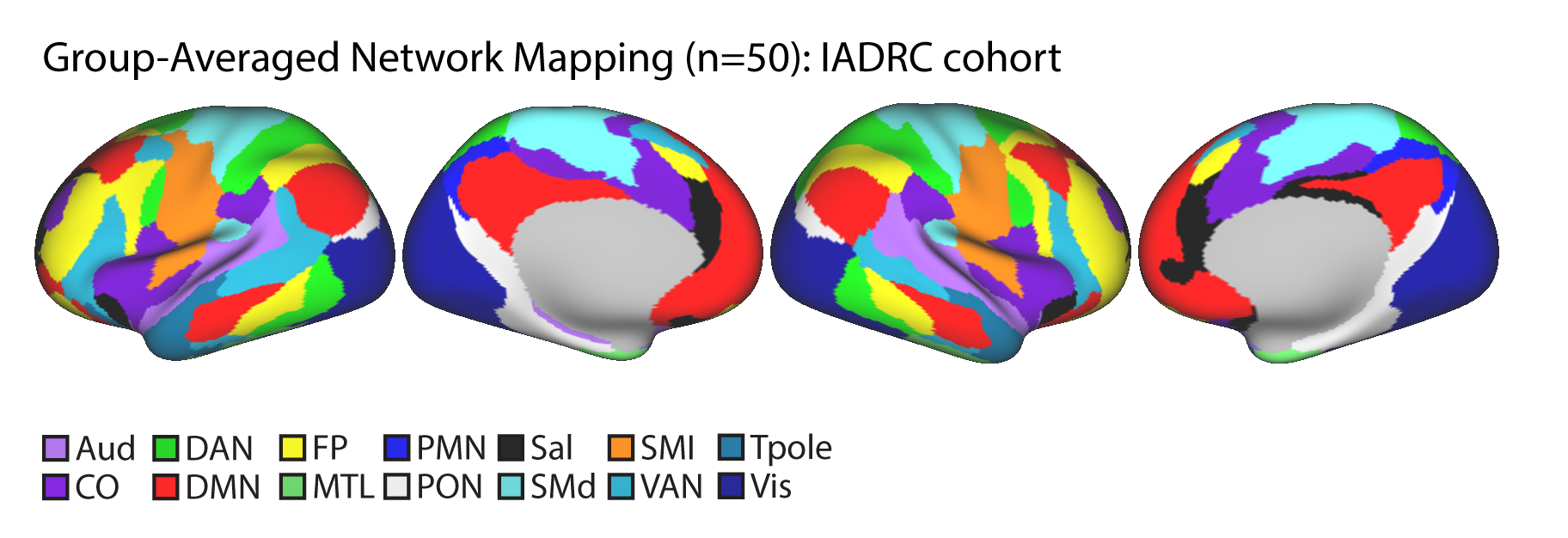

*Note*. This network map was created from the 50 (of 107) lowest motion IADRC participants. It is plotted on the HCP S1200 group-averaged very inflated surfaces.

**Supplemental Figure 6**

*Homogeneity of different cortical networks in the IU (a) and IADRC (b) cohorts using group-averaged versus individualized network mapping*

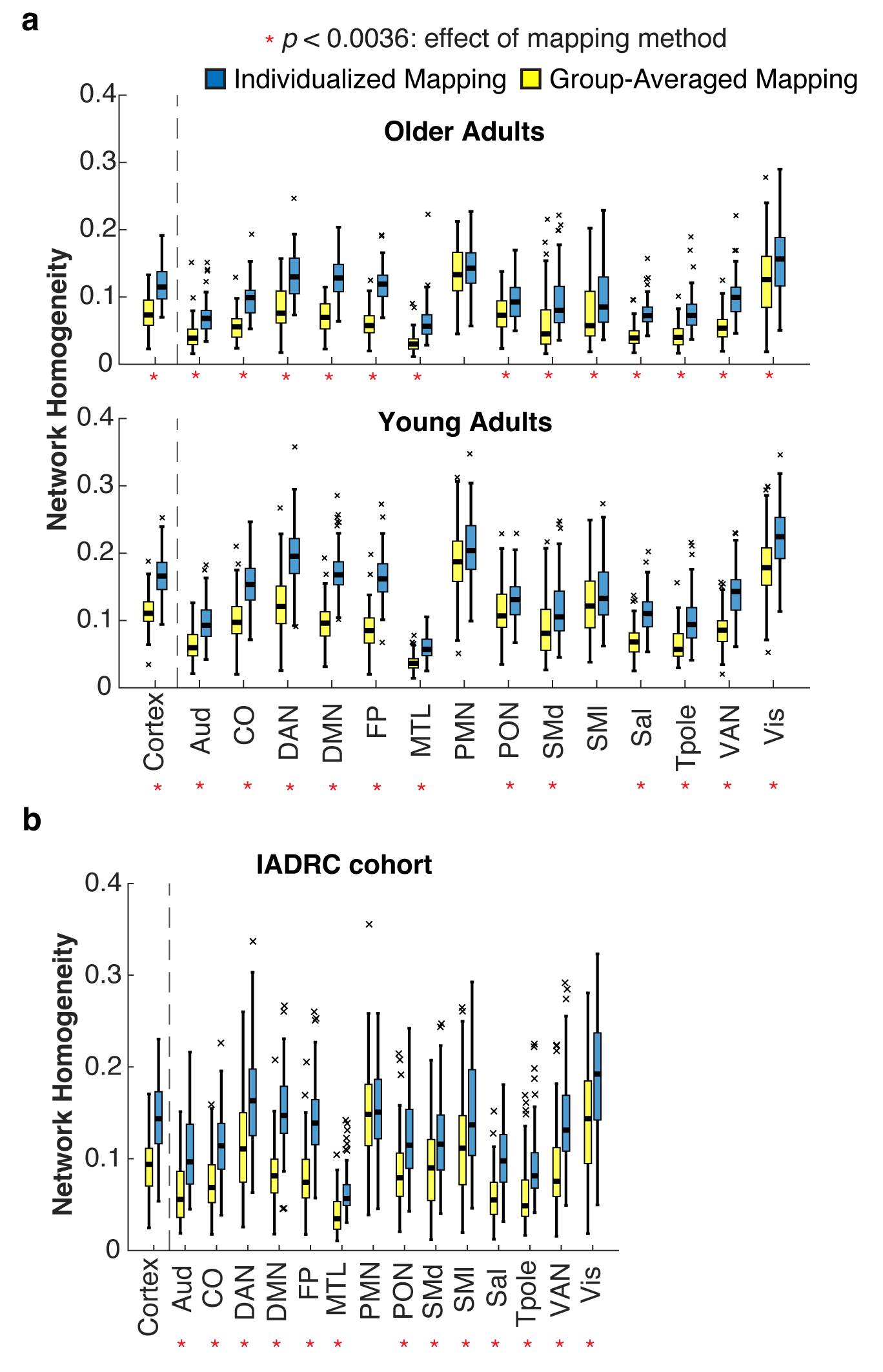

*Note.* Network homogeneity – the strength of the correlations between vertices within networks – is higher for individualized versus group-averaged network mapping. Networks that exhibited an effect of mapping method at the Bonferroni-corrected *p*-value of 0.0036 are noted with an asterisk.

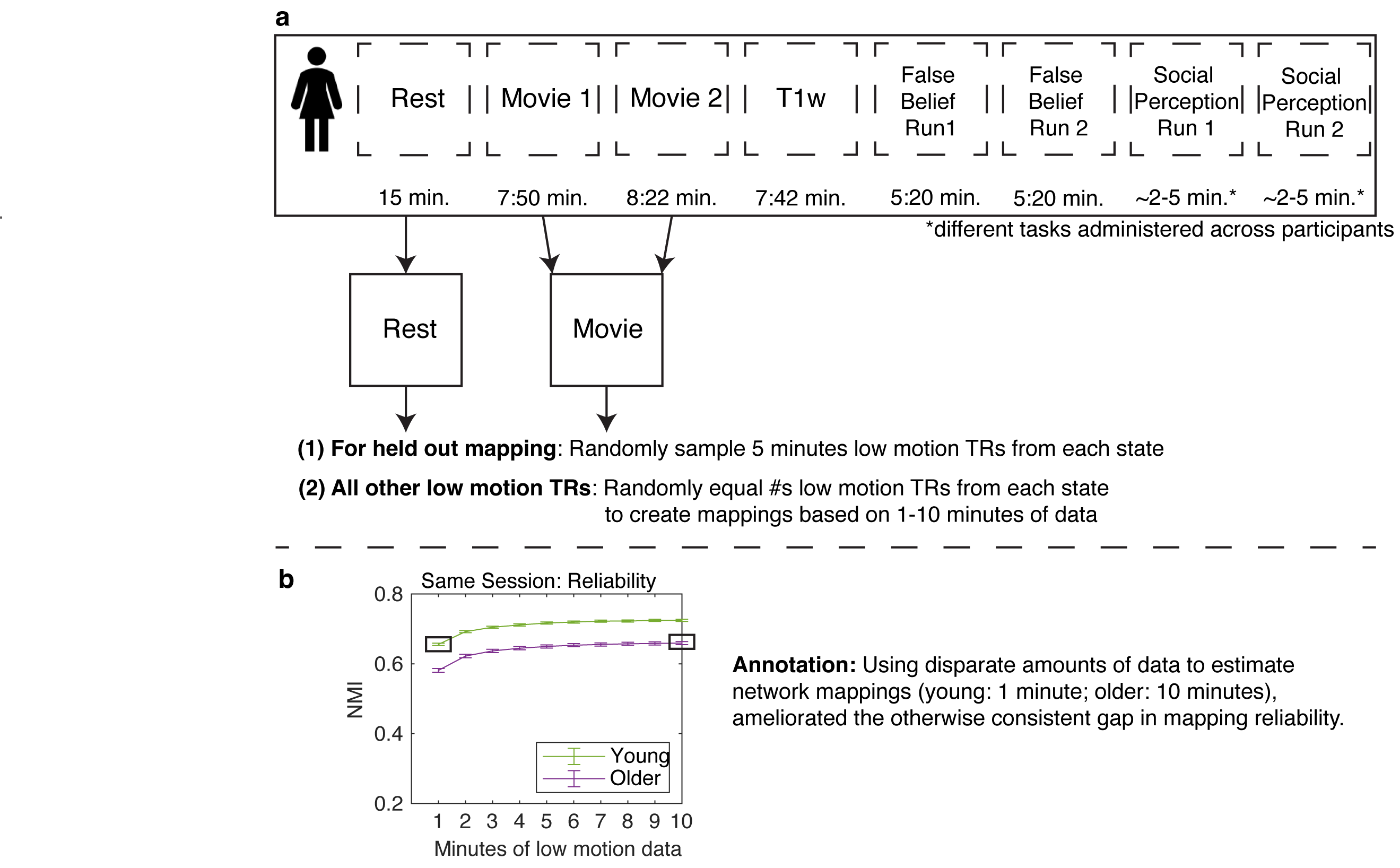
**Supplemental Figure 7**

**Supplemental Figure 8**

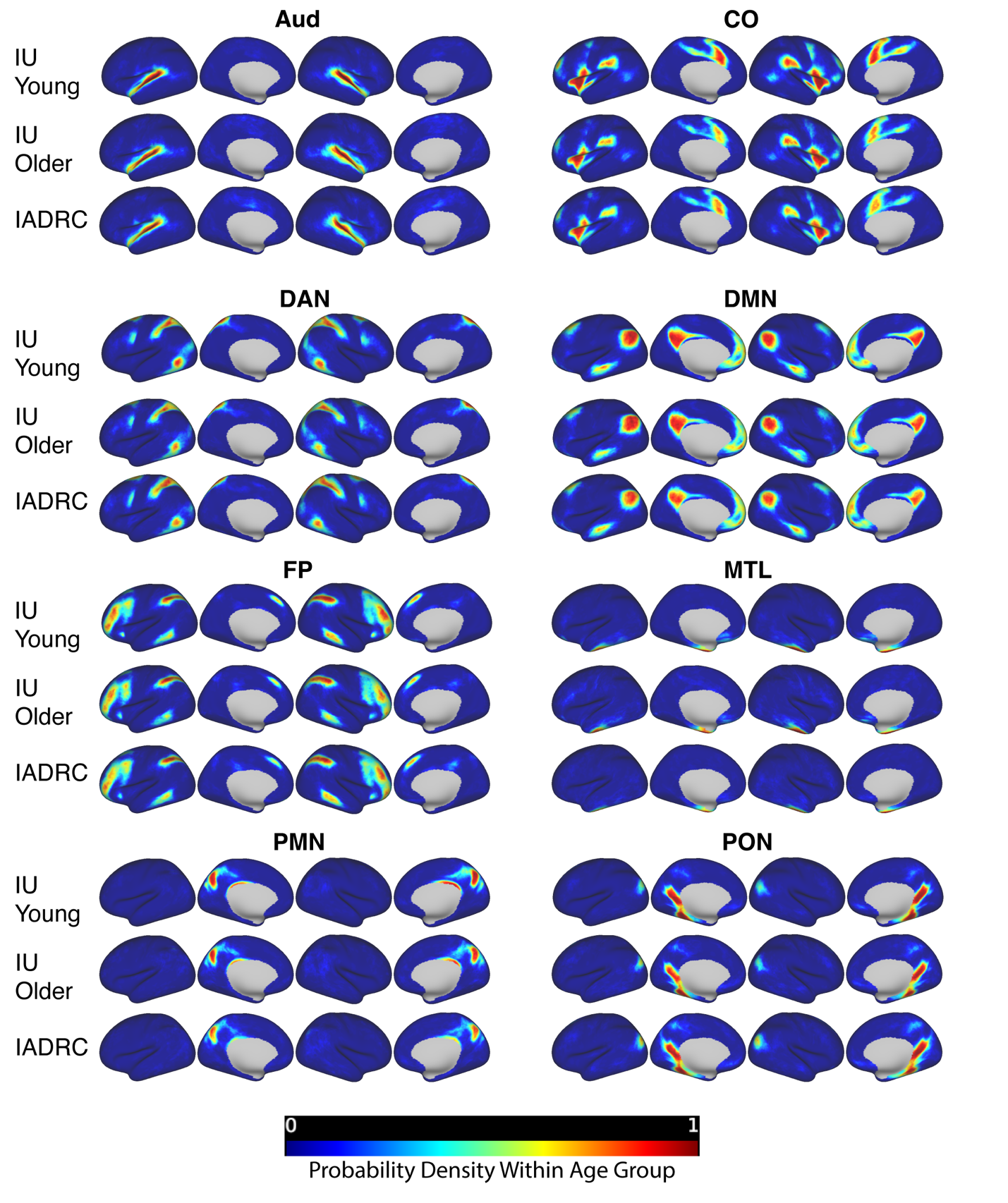
*Probabilistic Network Maps in IU and IADRC Cohorts*

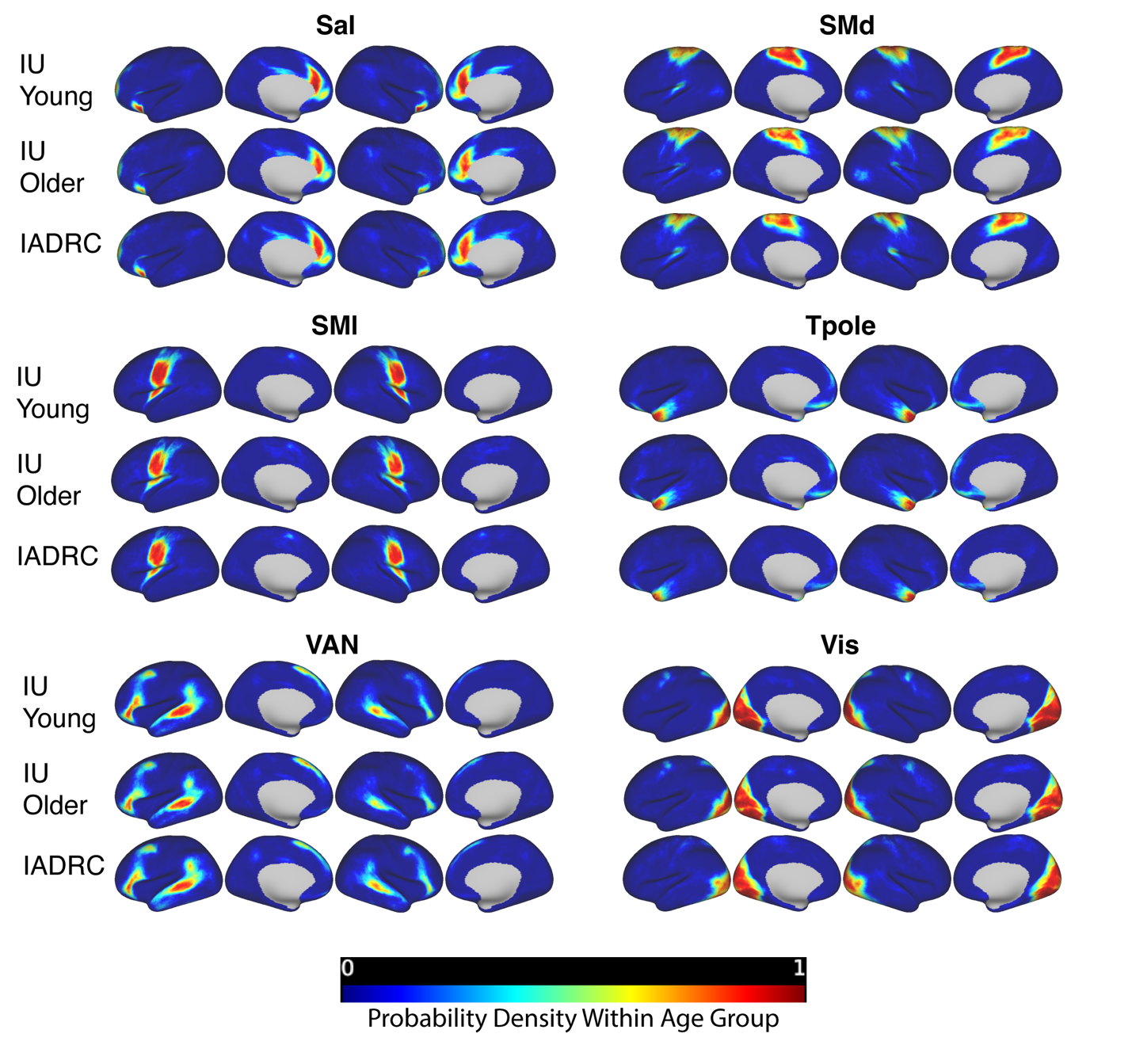

*Note*. Plot colors indicate the proportion of participants within each group for whom each vertex was assigned to the target network (i.e., probability density). Yellow is approximately 0.60 (60% probability) and orange is approximately 0.70 (70% probability).

**Supplemental Figure 9**

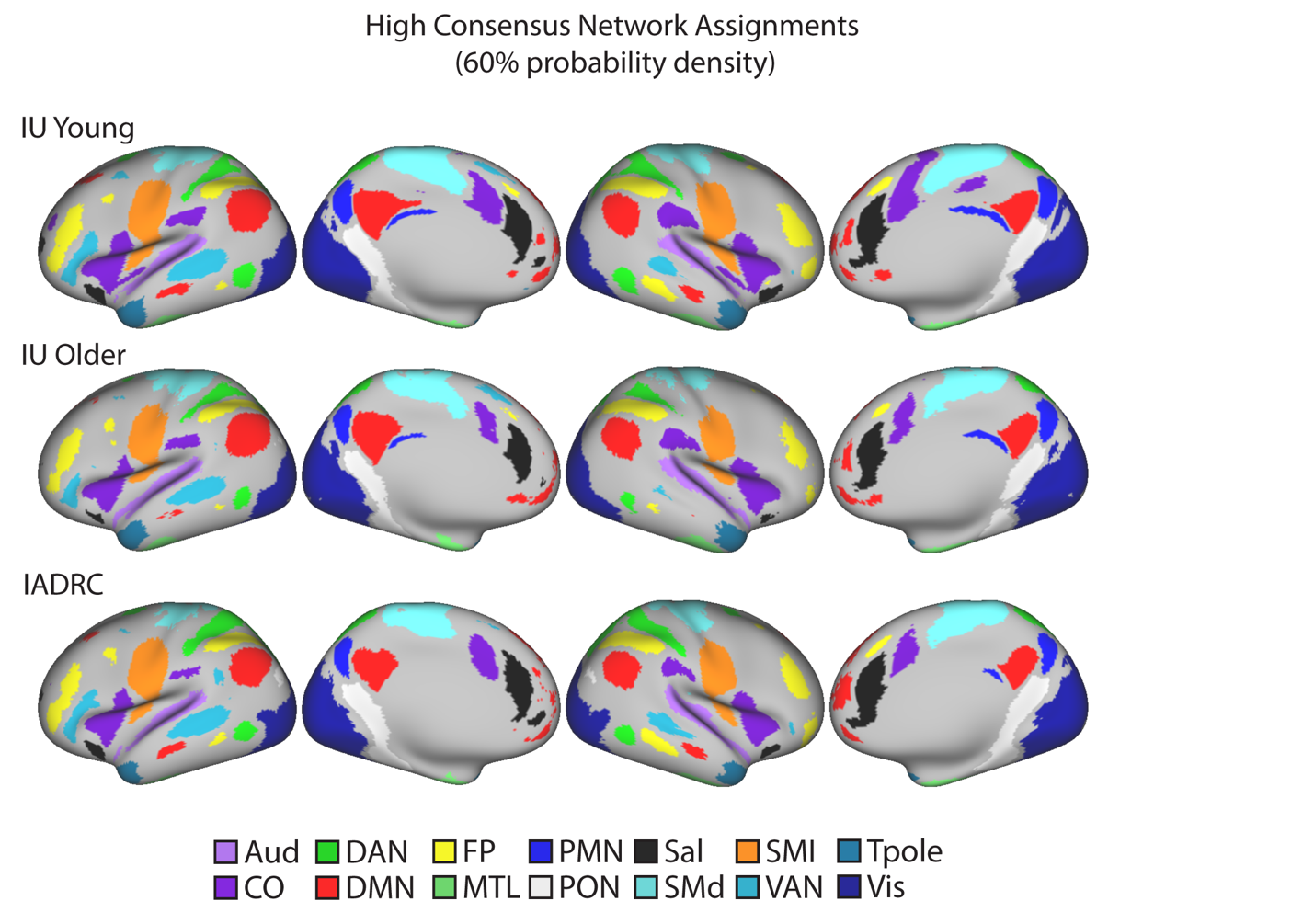
*High consensus network assignments using a 60% probability threshold*

*Note.* High probability network assignments in each group using an 60% probability threshold (see also Figure 4a which uses an 80% threshold).

**Supplemental Figure 10**

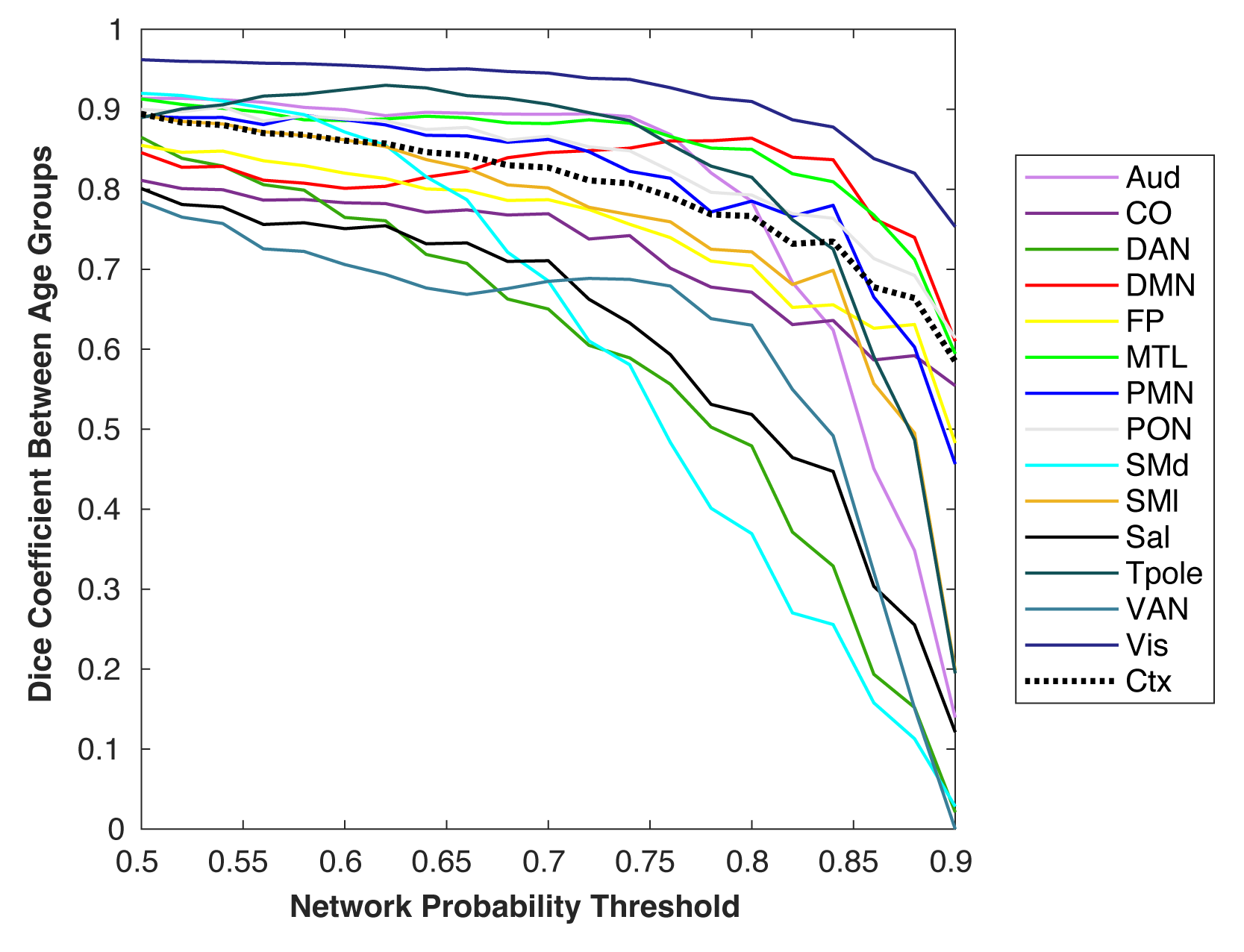
*Similarity between IU young and older adults in high probability network assignments across a range of network probability thresholds*

*Note.* We tested the similarity between high probability network assignments (e.g., Figure 4a, Supplemental Figure 9) across a range of network probability thresholds from 0.50 to 0.90 in increments of 0.02. The x-axis is simplified to increments of 0.05 for ease of visualization. The label ‘Ctx’ refers to the analysis of label consistency across networks.

**Supplemental Figure 11**

*Resting-state fMRI temporal signal-to-noise ratio (tSNR) maps for each group*
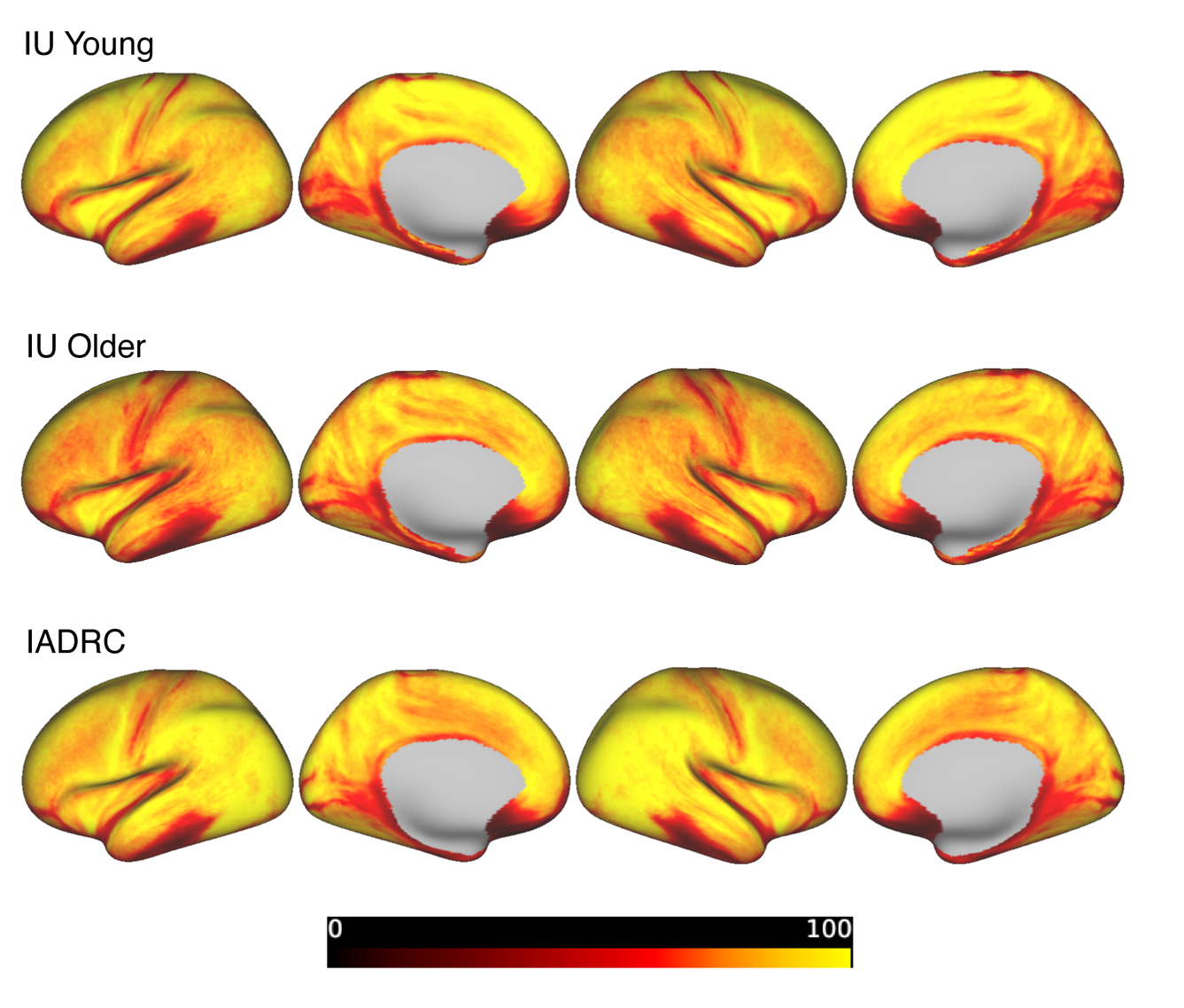

*Note*. The IU and IADRC cohorts were both acquired on a Siemens Prisma 3T MRI scanner but at different sites, using different head coils, and under different acquisition parameters (see *Methods*).
